# Quantitative analyses of interactions between SpoVG and RNA/DNA

**DOI:** 10.1101/2023.02.06.527361

**Authors:** Timothy C. Saylor, Christina R. Savage, Andrew C. Krusenstjerna, Nerina Jusufovic, Wolfram R. Zückert, Catherine A. Brissette, Md. Motaleb, Paula J. Schlax, Brian Stevenson

## Abstract

The *Borrelia burgdorferi* SpoVG protein has previously been found to be a DNA- and RNA-binding protein. To aid in the elucidation of ligand motifs, affinities for numerous RNAs, ssDNAs, and dsDNAs were measured and compared. The loci used in the study were *spoVG, glpFKD, erpAB, bb0242, flaB*, and *ospAB*, with particular focus on the untranslated 5’ portion of the mRNAs. Performing binding and competition assays yielded that the 5’ end of *spoVG* mRNA had the highest affinity while the lowest observed affinity was to the 5’ end of *flaB* mRNA. Mutagenesis studies of *spoVG* RNA and ssDNA sequences suggested that the formation of SpoVG-nucleic acid complexes are not entirely dependent on either sequence or structure. Additionally, exchanging uracil for thymine in ssDNAs did not affect protein-nucleic acid complex formation.

## 2. Introduction

SpoVG proteins are conserved across diverse groups of bacteria, including Firmicutes, Spirochaetota, and Myxococcota (Delta-proteobacteria) [1]. The protein’s nomenclature derives from early studies of *Bacillus* spp., where it was observed that mutations within the *spoVG* locus impaired sporulation at stage V [2]. The SpoVG homologues of *Borrelia burgdorferi* and other non-sporulating bacterial species affect expression of a broad range of genes, and *spoVG* mutants exhibit physiological defects [3–8]. Our group has studied SpoVG proteins from *B. burgdorferi, Staphylococcus aureus*, and *Listeria monocytogenes*, and found that they exhibit affinities for specific DNA sequences [1]. Subsequent studies by Burke and Portnoy found that *L. monocytogenes* SpoVG exhibits a substantially greater affinity for single-stranded RNAs than for cognate double-stranded DNAs (dsDNAs), and hypothesized that SpoVG functions primarily as an RNA-binding protein [3]. Evaluation of the *B. burgdorferi* homologue found a similarly greater affinity for RNAs [4].

The Lyme disease spirochete, *B. burgdorferi*, is a particularly useful model for studies of gene and protein regulation in a vector-borne pathogen [9–11]. In addition to being the cause of a significant human disease, *B. burgdorferi* infects both vertebrates and ticks [12]. The Lyme spirochete must, therefore, possess mechanisms to accurately determine which host it is in, and produce proteins and other essential factors for each stage of its infectious cycle. *B. burgdorferi* must also recognize when a tick is feeding, in order to undergo the physiological changes that are necessary for transmission into the vertebrate host. Our group’s previous data indicate that *B. burgdorferi* SpoVG affects expression levels of numerous proteins and affects bacterial physiology, apparently through its activities as an RNA- and/or DNA-binding protein [4]. To gain further insights into SpoVG function, we identified high- and low-affinity ligands of *B. burgdorferi* SpoVG, which were then employed to gain insights on the ability of SpoVG to bind single-stranded and double-stranded nucleic acids.

## 3. Materials and Methods

### 3.1. Purification of recombinant proteins

Polyhistidine-tagged *B. burgdorferi* SpoVG was purified essentially as described previously [1,4]. Briefly, *Escherichia coli* Rosetta II (Invitrogen, MA) was transformed with pBLJ132, which consists of *spoVG* cloned into pET101 [1]. *E. coli* were then grown to an OD_600_ of at least 1.0 in Super Broth (32 g Tryptone, 20 g Yeast Extract, and 5 g NaCl per liter), and recombinant SpoVG expression was induced for 1h by adding isopropyl-β-D-thiogalactopyranoside (IPTG) to a final concentration of 1mM. Bacteria were harvested by centrifugation at 5400 x *g* for 30 minutes and frozen at −80°C until needed. Resuspended cells were lysed by sonication with the addition of B-PER bacterial protein extraction reagent to 2% v/v (Thermo-Fisher, MA). Recombinant proteins were purified using MagneHis nickel particles (Promega, WI), then dialyzed against EMSA buffer (50mM Tris-HCl, 25mMKCl, 10% glycerol (v/v), 0.01% Tween 20, 100nM dithiothreitol (DTT), and 1mM phenylmethanesulfonyl fluoride (PMSF)). Proteins were concentrated using 10kDa Amicon centrifugal units (MilliporeSigma, MA) and aliquots were stored at −80°C until needed. Protein purity and concentration were assessed by SDS-PAGE, Quick Start Bradford protein assay (Bio-Rad), and bicinchoninic acid assay (BCA) (Thermo-Fisher, MA).

### 3.2. Electromobility shift assay (EMSA)

Fluorescently tagged and untagged DNA and RNA oligonucleotides were synthesized by Integrated DNA Technologies (IDT, IA). The sequences of oligonucleotides used in this study are listed in Table 1 and Supplemental Table 1. DNA and RNA probes were tagged on the 5’ end with an IRDye 800 fluorescent tag (LI-Cor, NE) or an Alexa Fluor 488, respectively. Double-stranded DNA (dsDNA) probes and competitors were produced from pairs of separately-synthesized complimentary oligonucleotides by mixing equal molar concentrations, heating to 95°C, and slowly cooling to room temperature.

**Table 1.**
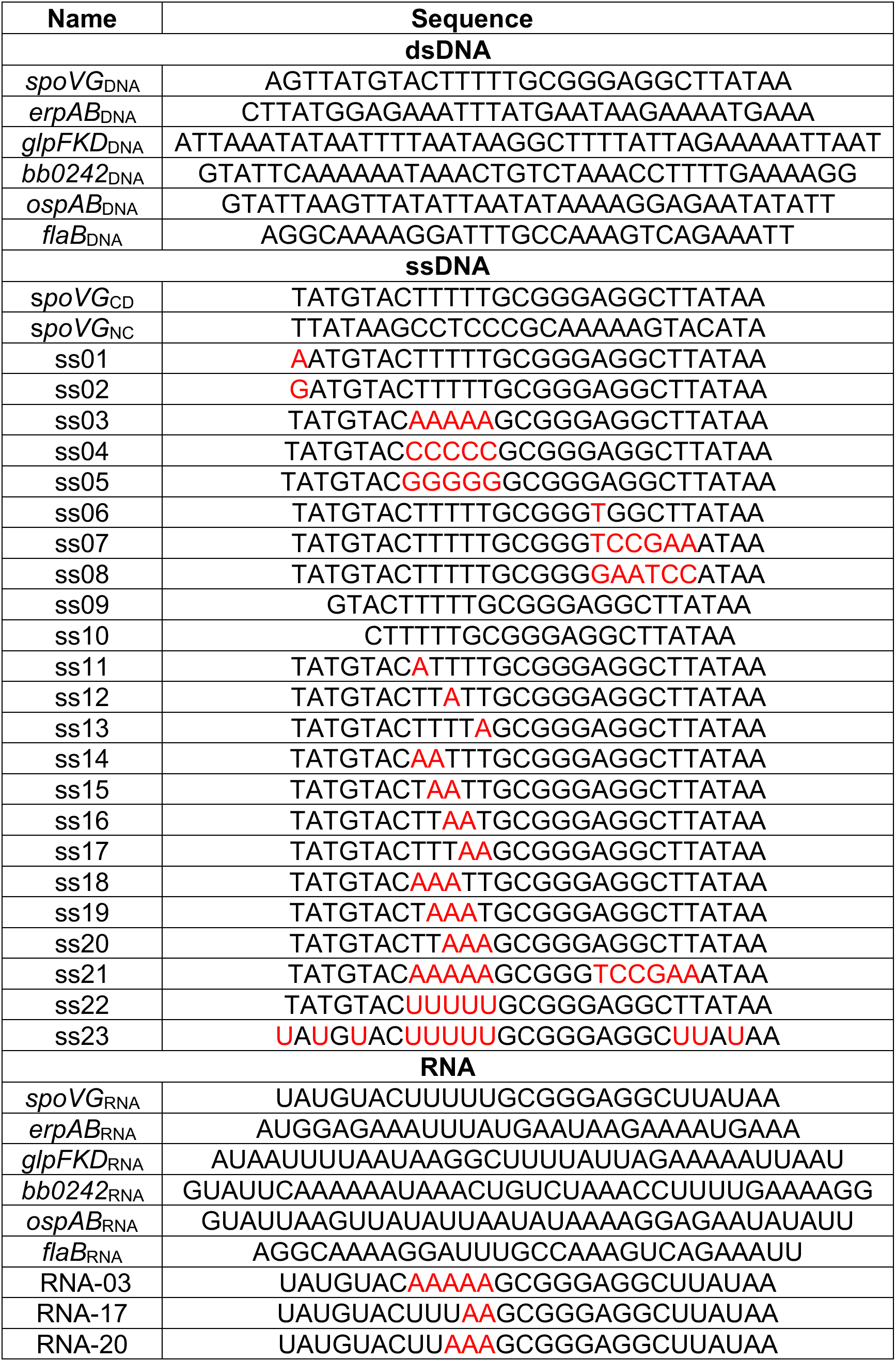
5’-3’ sequences of dsDNA, ssDNA and RNA used throughout. Red highlighted nucleotides in the ssDNA and RNA sections deviate from the *spoVG* probe. ss22 and ss23 have a deoxyribose backbone with uracil bases instead thymine.

EMSAs were performed as previously described [1,4]. Unless stated otherwise, purified proteins were added to 100nM of labeled nucleic acid probes and incubated at room temperature for 5-15 minutes. Unlabeled nucleic acid competitors were added prior to the addition of protein. EGTA was added to the indicated reactions to inhibit RNAse activity in protein aliquots. Poly-dI-dC was added to indicated gels as a non-specific competitor. One-sixth volume of EMSA loading dye (15mg/mL Ficol 400, 0.8 mg/mL Orange G) was added. Electrophoresis was performed using pre-run 6% TBE gels (Invitrogen, MA) at 100-volts in 0.5x TBE buffer. Gels were imaged and analyzed via densitometry with a ChemiDoc MP and Image-Lab software, respectively (Bio-Rad, CA). Lanes and bands were added manually according to the strongest free probe and the strongest shifted band. Equivalent bands were then added across the entire EMSA.’ Background noise was calculated and accounted for by the Image-Lab software. Percent shifted was graphed using GraphPad Prism 9 (Dotmatics, MA).

### 3.3. Calculation of apparent K_D_ and IC50 values

For direct binding assays, the fraction of bound complex was fit to a one-site saturation ligand binding model (Fraction Bound= Bmax SpoVG/ (Kd + SpoVG)) where Bmax is the maximum fraction bound and K_D_ is the apparent dissociation equilibrium constant [13]. It should be noted that this estimate of the binding constant is limited by the high concentration of nucleic acid in the binding assays which likely puts acids in the “Titration Regime” but can still be used to determine relative affinities of one sequence to another [13]. In competition assays, data were fit globally to determine the concentration of inhibitor resulting in 50% of the complex being disrupted (IC50) to the equation fraction bound to labeled probe = m – (m-b)/(1+IC50/[C]) where m is the maximum signal, b is the minimum (background) signal, [C] is the concentration of the unlabeled nucleic acid competitor, and IC50 is the concentration of the competitor that results in 50% of the complex being disrupted.

## 4. Results

### 4.1. SpoVG is a site-specific RNA- and DNA-binding protein

We previously reported that *B. burgdorferi* SpoVG can bind to the 5’ ends of the borrelial *spoVG* and *glpFKD* mRNAs, and then cognate dsDNAs [4]. It was also found that SpoVG binds within the transcript of a small gene of unknown function, *bb0242*, that is located between the *glpK* and *glpD* genes [4]. The present studies delved further into the interactions between SpoVG and other nucleic acid sequences to assess relative affinities and mechanisms of binding. Noting that *spoVG* and the *glpFKD* operon are highly expressed during tick colonization but not during mammalian infection, we investigated the ability of SpoVG to bind other borrelial sequences; specifically the 5’ ends of *erpAB* (which is repressed during tick colonization but highly expressed during mammalian infection), *ospAB* (which is highly expressed during tick colonization but repressed in mammals), and *flaB* (which is constitutively expressed throughout the borrelial tick-mammal cycle) [4,12,14–18]. Both RNA and dsDNA were examined for each target.

Of the tested probes and competitors, SpoVG exhibited the greatest affinity for the *spoVG* mRNA 5’ end, henceforth referred to as *spoVG*_RNA_ (Fig. 1 and Fig. 2). This high affinity binding site was further tested by competition EMSAs using an unlabeled *spoVG*_RNA_ as a competitor (Fig. 1A and Supp. Fig. 1). No intermediate shift was observed indicating that there was no formation of a dsRNA probe. The protein-RNA complex shift stayed above 50% until 1μM (10x) of competitor was added. Addition of up to 100 ng/μL poly-dI-dC did not result in an appreciable difference of SpoVG binding to the *spoVG*_RNA_ (Fig. 1B), further indicating that binding of SpoVG to *spoVG*_RNA_ is specific. SpoVG also bound to the *glpFKD*_RNA_, *bb0242*_RNA_, and *erpAB*_RNA_ probes, but EMSAs did not approach saturation in the tested ranges of SpoVG protein concentrations, indicating that the affinity of SpoVG for those three RNAs is weaker than *spoVG*_RNA_ (Fig. 2).

**Figure 1.**
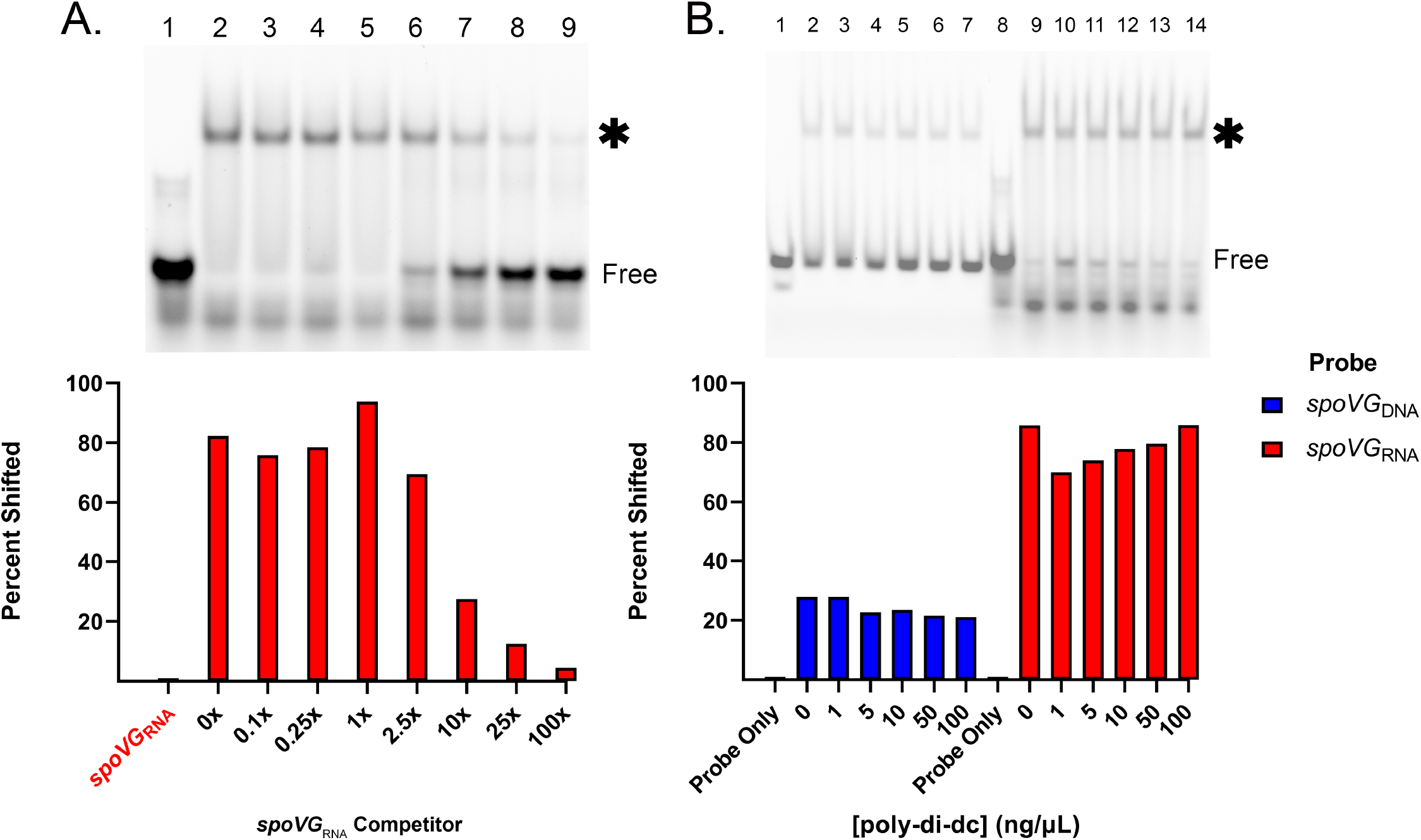
*spoVG* RNA is a SpoVG high affinity binding site. **A.** Competition assay using 100nM labeled *spoVG*_RNA_. Free probe is indicated, and the shifted probe is indicated by an asterisk. SpoVG concentration was 500nM in lanes 2-9. Unlabeled *spoVG*_RNA_ was added at 10nM, 25nM, 100nM, 250nM, 1μM, 2.5 μM, and 10.0 μM in lanes 3-9 respectively. EGTA was added to 5mM. This assay was done in triplicate (Sup. Fig. 1). **B.** Competition assay using labeled *spoVG*_DNA_ and *spoVG*_RNA_ probe against poly-dI-dC. Lanes 1-7 included 100nM *spoVG*_DNA_ probe. Lanes 8-14 included 100nM *spoVG*_RNA_ probe. SpoVG was added to lanes 2-7 at 10μM and lanes 9-14 at 100nM. Poly-dI-dC was added in at 1,5,10,50, and 100 ng/μL in lanes 3-7 and 10-14 respectively.

**Figure 2.**
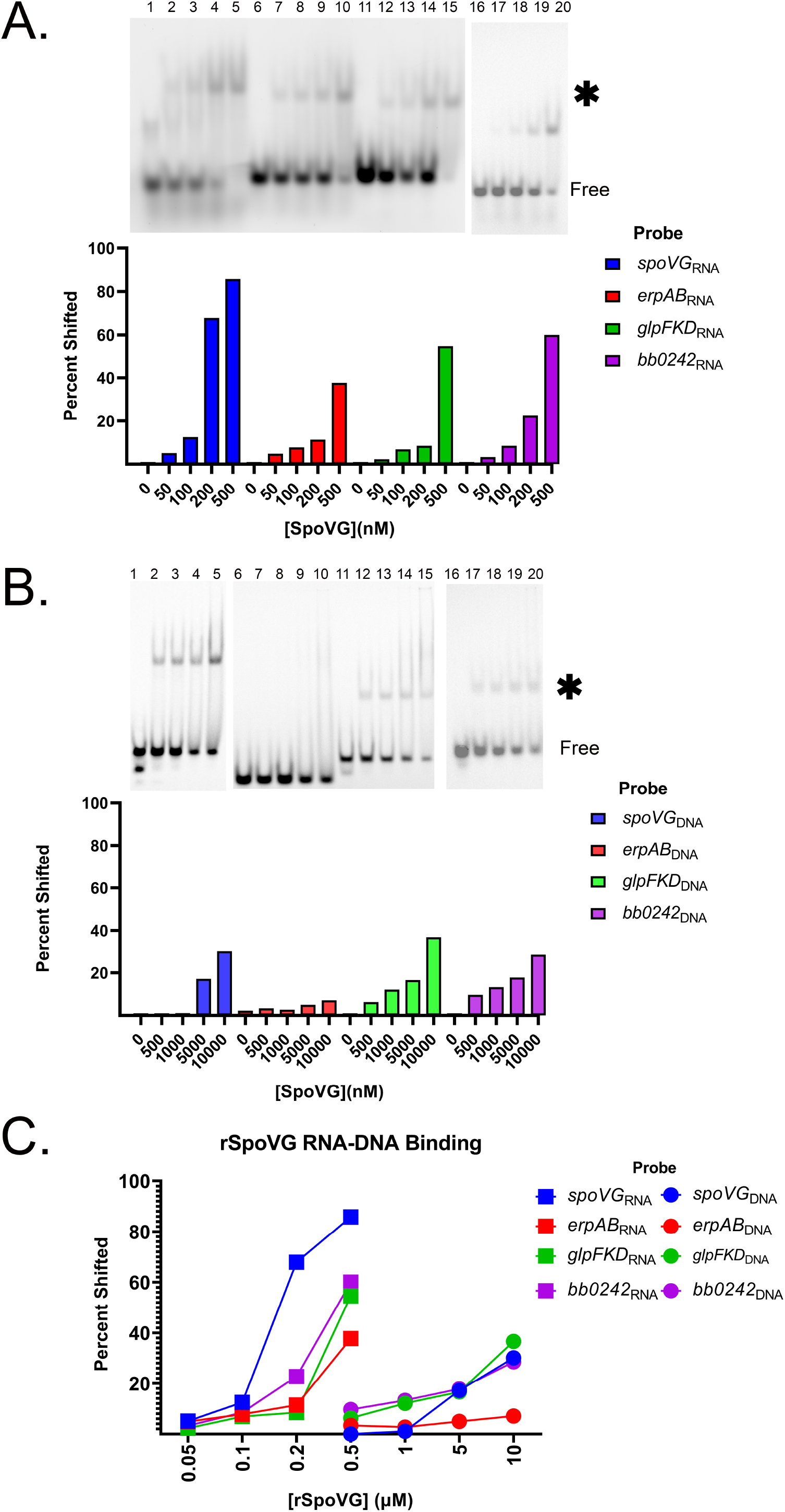
SpoVG has a higher binding affinity for RNA than DNA. **A.** EMSA using labeled *spoVG*_RNA_, *erpAB*_RNA_, *glpFKD*_RNA_, and *bb0242*_RNA_ at 100nM from lanes 1-5, 6-10, and 11-15, 16-20 respectively. Free probe is indicated, and the shifted probe is indicated by an asterisk. SpoVG was added to lanes 2-5, 7-10, 12-15, and 17-20 at 50nM, 100nM, 200nM, and 500nM. **B.** EMSAs using labeled *spoVG*_DNA_, *glpFKD*_DNA_, *erpAB*_DNA_, and *bb0242*_DNA_ at 100nM from 1-5, 6-10, 11-15, 16-20 respectively. SpoVG was added to lanes 2-5, 7-10, 12-15, and 17-20 at 500nM, 1μM, 5 μM, and 10 μM respectively. **C.** Graph representing the percentage of 100nM probe shifted due to SpoVG binding from the EMSAs in A and B.

Our initial studies of *B. burgdorferi* and *S. aureus* SpoVG found that both proteins preferentially bound to certain dsDNA sequences [1]. In contrast, a study of the *L. monocytogenes* orthologue with a variety of DNA probes led those researchers to conclude that DNA binding appeared to be non-specific [3]. To build upon those observations, we assessed the affinities of *B. burgdorferi* SpoVG for the above-described borrelial sequences as dsDNA. Affinities for all tested dsDNAs were significantly lower than their cognate RNAs (Fig. 2). The relative affinity of SpoVG to the *spoVG*_DNA_ probe was over 100x weaker than for *spoVG*_RNA_, with addition of 10μM SpoVG protein yielding approximately 30% shifted probe (Fig. 2B). This difference signifies that a substantially higher concentration of SpoVG is required to form a complex with SpoVG_DNA_ dsDNA compared to SpoVG_RNA_.

These results were highlighted by performing EMSAs of the *spoVG*_DNA_, *spoVG*_RNA_, *glpFKD*_DNA_, and *glpFKD*_RNA_ probes (Supp. Fig. 2). EMSAs with *spoVG* RNA achieved close to 100% shift at 200nM compared to the unshifted dsDNA at 200nM concentration (Supp. Fig. 3A). This was also observed with the *glpFKD*_RNA_. Saturation of the *glpFKD*_RNA_ probe occurred between 1μM and 10μM SpoVG, whereas the *glpFKD*_DNA_ probe showed almost no detectable shift at that concentration. While SpoVG had an inherently weaker interaction with the DNAs tested than the RNAs, there were still substantial differences when comparing affinities for the DNA probes out of the four tested. SpoVG had the lowest relative affinity for the *erpAB*_DNA_ probe (Fig. 2B and C).

The relative affinities of SpoVG for the *ospAB*_RNA_ and *flaB*_RNA_ were determined by use of unlabeled RNAs as competitors against labeled *spoVG*_RNA_ or *glpFKD*_RNA_ probes. When using the *spoVG*_RNA_ probe, addition of 100x excess of the unlabeled *ospAB*_RNA_ reduced the shifted protein-RNA complex by approximately 18% (IC50 = 5.9 +/- 4.6 × 10^−5^ M), while 100x excess of the unlabeled *flaB*_RNA_ did not detectably affect the SpoVG-*spoVG*_RNA_ complex (Fig. 3A and B). When labeled *glpFKD*_RNA_ was used as a probe, addition of 25x excess of unlabeled *flaB*_RNA_ led to a 20% reduction in the SpoVG-*glpFKD*_RNA_ complex (Fig. 3C). In contrast, 25x excess of unlabeled *glpFKD*_RNA_, *erpAB*_RNA_, or *ospAB*_RNA_ reduced binding to the probe by approximately 80%, and unlabeled *spoVG*_RNA_ virtually eliminated the SpoVG-*glpFKD*_RNA_ complex (Fig. 3C). The data above indicate that SpoVG has a substantially greater affinity for RNA compared to DNA, but that this affinity differs based on the sequence of the nucleic acid.

**Figure 3.**
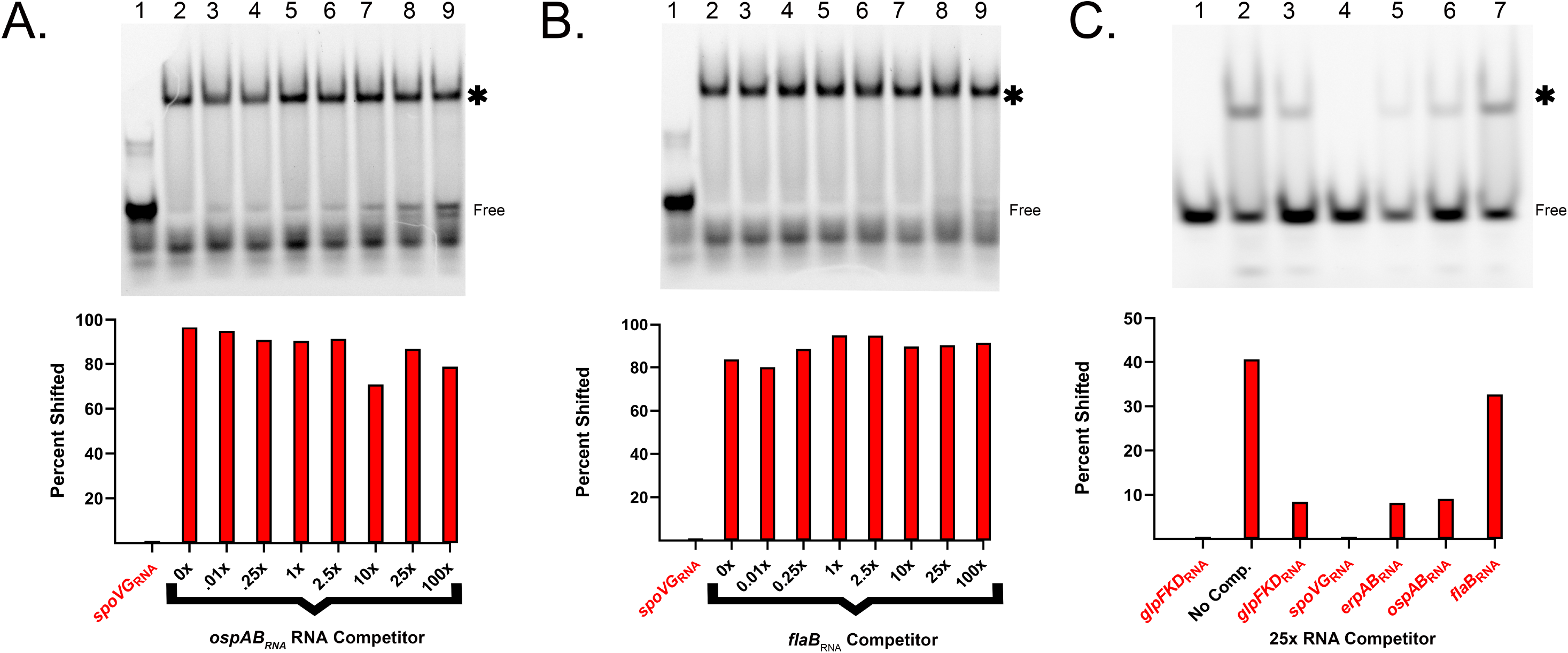
SpoVG is a specific RNA binding protein. **A.** Competition assay using 100nM labeled *spoVG*_RNA_. Free probe is indicated, and the shifted probe is indicated by an asterisk. The SpoVG concentration was 500nM in lanes 2-9. Unlabeled *ospAB*_RNA_ was added at 10nM, 25nM, 100nM, 250nM, 1μM, 2.5 μM, and 10.0 μM in lanes 3-9 respectively. EGTA was added to 5mM. **B.** Competition assay using unlabeled *flaB*_RNA_ and 100nM labeled *spoVG*_RNA_. Lane values match (A), except unlabeled *flaB*_RNA_ was used instead of *ospAB_RNA_*. **C.** Competition assay using 100nM labeled *glpFKD*_RNA_ (lanes 8-14). SpoVG concentration was 500nM. Competitors were added at a concentration of 2.5μM in the following sequence *glpFKD*_RNA_, *spoVG*_RNA_, *erpAB*_RNA_, *bb0242*_RNA_, and *flaB*_RNA_. The gamma of entire image was increased to more accurately visualize differences in competition.

### 4.2. Effects of nucleic acid sequence on SpoVG-binding

As the SpoVG protein appears to have the strongest affinity for the *spoVG_RNA_* probe, we made a series of sequence variants to determine whether the sequence or structure of the RNA was important in recognition and affinity of the protein to the nucleic acid. EMSAs were undertaken using the labeled *spoVG*_RNA_ probe and numerous unlabeled variant nucleic acids as competitors (Table 1). As a cost saving first approach, ssDNA competitors were used for the initial analyses. Sequence variations in competitors included changing the 5’ nucleotide, removing 3 or 6 nucleotides from the 5’ end, altering the run of five consecutive thymines, and altering an AGGCT sequence near the 3’ end (Table 1).

Of the 21 tested unlabeled ssDNAs, only a subset of the oligonucleotides that disrupted the five-thymine run had appreciable impacts on competition for SpoVG (Fig. 4A and B). Competition of the *spoVG*_RNA_ with nucleic acid fragments that replace the sequence TTTTT in the unlabeled probe with CCCCC (ss04) or GGGGG (ss05) did not appreciably affect their ability to compete with the wild type *spoVG* sequence (Fig. 4A). However, replacing the run of thymines in the sequence with adenines in some positions: (AAAAA) (ss03), TTTAA (ss17), or AAATT (SS18), or with Us (ss21) resulted in reduced competition. Lastly, when combining the changes of the TTTTT sequence to AAAAA with deletion of the AGGCT sequence (ss21), the competition was reduced but was similar to that of the AAAAA (ss03) change only.

**Figure 4.**
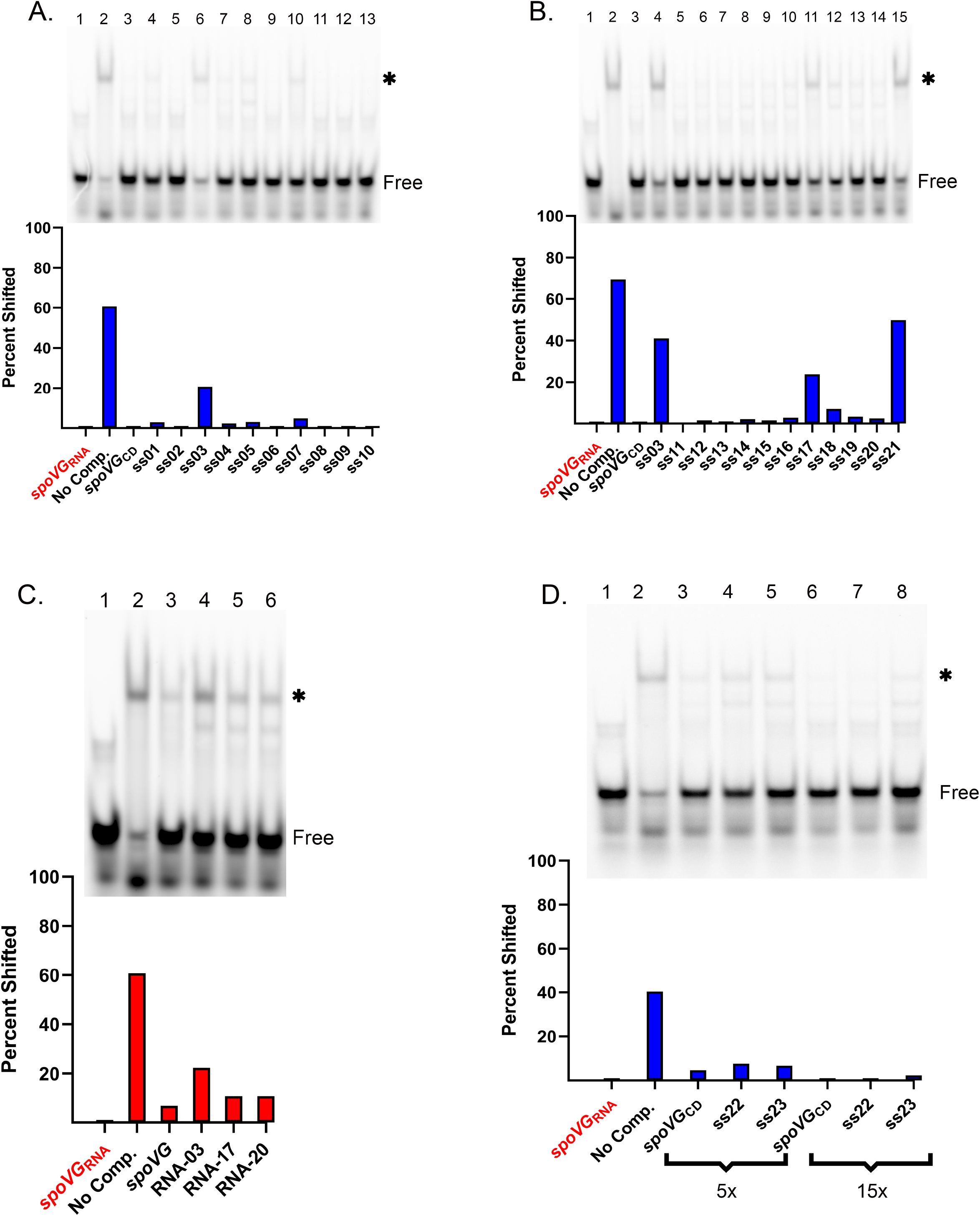
Mutagenesis of *spoVG* ssDNA and RNA. **A.** EMSA using 100nM *spoVG*_RNA_ probe and 100nM of SpoVG in lanes 2-13. ssDNA competitors were added to lanes 3-13 at 5μM. Free probe is indicated, and the shifted probe is indicated by an asterisk. Competitors used were *spoVG+* through ss10. (Table 1 and Supp. Table 1) **B.** EMSA using 100nM *spoVG*_RNA_ probe and 100nM of SpoVG in lanes 2-15. Competitors at 5μM were added to lanes 3-15. Competitors were *spoVG*_CD_, ss03, followed by ss11 through ss21. (Table 2) **C.** EMSA using 100nM *spoVG*_RNA_ probe with 100nM SpoVG added to lanes 2-6. Unlabeled *spoVG*_RNA_, RNA-03, RNA-17, and RNA-20 were added at 1.5μM to lanes 3-6 respectively. **D.** EMSA using 100nM of *spoVG*_RNA_ probe and 100nM of SpoVG added to lanes 2-8. ssDNA competitors consisted of *spoVG*_CD_, ss22, and ss23 at 500nM in lanes 3-5 and 2μM in lanes 6-8.

Based on above results, three unlabeled RNAs were synthesized for use as EMSA competitors: RNA-03 changed the run of five uracils (UUUUU) to adenines (AAAAA), RNA-17 changed the run to UUUAA, and RNA-20 changed it to UUAAA (Table 1). When added to EMSAs at 15-fold excess, each of these mutant competitors was slightly less effective at competing for SpoVG than the spoVG_RNA_ sequence (Fig. 4C). RNA-03 saw a 29% decrease in competition, whereas RNA-17 and RNA-20 saw a 7% decrease in competition compared to the wild type sequence. All RNAs effectively competed for SpoVG when added at 50x excess whereas the ssDNA counterparts, ss03 and ss17, were less effective than *spoVG*_CD_ (Table 1 and Fig. Supp. 3A). In contrast, 50x excess of unlabeled *glpFKD*_RNA_ competitor did not compete nearly as well as any tested *spoVG* mutant RNA sequence. (Fig. Supp. 3B). Altogether, this suggests that SpoVG differentially interacts with additional aspects of the *spoVG*_RNA_ that are not present in the *glpFKD*_RNA_.

Due to the observed differential affinities above between the RNA probes derived from the start of transcription and their cognate dsDNAs, we performed competition assays comparing the coding (*spoVG*_CD_) and non-coding (*spoVG*_NC_) strands of DNA. EMSAs were performed in which the two ssDNAs competed against labeled *erpAB*_RNA_, which was used to inhibit possible base pairing. This competition EMSA indicated that *spoVG*_NC_ better competed against *erpAB*_RNA_ than *spoVG*_CD_ (Supp. Fig. 4A and B). A substantial difference was observed when competitors were added at 100nM concentration: *spoVG*_NC_ was able to compete away 62% of the SpoVG-*erpAB*_RNA_ complex compared to *spoVG*_CD_, which only competed away 12%.

### 4.3. Differences of Uracil vs Thymine alone can not explain the differences in RNA and ssDNA binding

The differential competition of the SpoVG derived ssDNA and RNA sequences lead us to further investigate whether the presence of uracils in RNA and thymines in ssDNA contributed to differences in SpoVG affinity. To that end, we produced two ssDNAs based on the *spoVG*_CD_ sequence: competitor ss22 replaced only the TTTTT with UUUUU, whereas competitor ss23 replaced every thymine in the sequence with uracil (Table 1). Competition analysis using the altered ssDNA sequences did not result in more effective competition than the original SpoVG ssDNA sequence at 5x and 15x competitor concentrations (Fig. 4D).

## 5. Discussion

*B. burgdorferi* SpoVG is a sequence-specific RNA- and DNA-binding protein, exhibiting binding constants of 5.9 +/- 2.1 x10^−8^ M from direct binding assays and with an apparent IC50 of 1.033 × 10^−6^ M +/- 2.7 × 10^−7^ for *spoVG*_RNA_. Binding affinities for tested dsDNAs appear to be over 100-times weaker than for cognate RNAs, yet selectivity and differential affinity for certain dsDNA sequences was observed.

Analyses with the high-affinity RNA sequence derived from the *spoVG* start of transcription provided insights on the nature of those interactions. This RNA contains a run of five uracils, UUUUU, which, when changed to adenines, AAAAA, substantially affected SpoVG binding. Two additional base replacements, UUUUU to UUUAA or UUAAA, also led to decreases in SpoVG-RNA complex formation. Note that two other RNAs that SpoVG bound with high affinity also contains runs of uracils: the *glpFKD*_RNA_ probe contains two poly-U tracts, UUUUAAU and UUUUAUU, and the *erpAB*_RNA_ probe contains one poly-U tract, UUUAU. However, *ospAB*_RNA_ contains runs of only two sequential uracils, UU, and the *flaB*_RNA_ contains a single uracil tract of UUU. These results suggest that a poly-U tract may play a role in SpoVG binding. Noting that unlabeled *glpFKD*_RNA_ was less effective than any of the mutated *spoVG* RNAs, it is evident that additional features have important impacts on complex formation.

Having observed that *spoVG* ssDNA and RNA sequence variants showed a wide variation in their abilities to bind SpoVG, we used RNA structure v6.4 to predict the secondary structures of these sequences and the other natural substrates that were observed (Supp. Table 1) [19]. A majority of the constructs that were the least effective competitors e.g. (SS-03 and RNA-03, SS-17 and RNA-17, SS-20 and RNA-20) are predicted to share a secondary structure where the nucleic acid has a stem of 4 or 5 base pairs and a loop with a sequence GGAG. Lacking a 2’ hydroxyl group, the RNA sequences may fold into a more A-form helix compared to the same sequence of DNA, providing a possible rationale for why SpoVG prefers RNA over DNA. This seems particularly likely in that replacing thymines with uracils (ss22 and ss23) was insufficient to increase affinity of the SpoVG protein for these sequences. Yet, sequence ss21, which is predicted to have a different structure, with variations in both the stem and the loop, was still an efficient competitor, suggesting that there are likely other determinants to binding.

The greater affinity of SpoVG for RNA over cognate ssDNA containing uracil implies that SpoVG is more likely to form complexes with RNA rather than DNA either through direct interactions with the backbone or because of RNA’s preferred tertiary folds compared to those of DNA. Previous site-directed mutagenesis of SpoVG revealed that two domains are involved with nucleic acid-binding [1]. Residues in the sole alpha helix of the protein confer nucleic acid sequence preference. Two positively charged residues (arginine or lysine) in a separate loop are conserved across all bacterial species and changing either residue to an alanine eliminated nucleic acid binding. Taken together, these results suggest that the charged residues of the loop may bind to the sugar-phosphate backbone and may be better positioned to interact with ribose rather than deoxyribose.

We acknowledge that the sequences used in this study were short fragments of the 5’ end of mRNAs and lack the complexity of full-length mRNAs of coding genes. We are presently examining the *B. burgdorferi* transcriptome by RNA immunoprecipitation - sequencing (RIP-Seq) to identify additional high-affinity SpoVG binding sites, which can then be compared to the binding sites herein, to provide an enhanced rational approach to characterizing RNA features that are involved with SpoVG-binding.

In conclusion, the results of these studies indicate that there is definitive preferential affinity of SpoVG for certain RNAs and DNAs within the *B. burgdorferi* genome. Furthermore, SpoVG can bind to numerous sites throughout the transcriptome and genome. *B. burgdorferi* controls levels of SpoVG throughout the infection cycle [4]. Thus, differential expression of SpoVG will result in occupancy of greater or fewer binding sites, facilitating a variety of different phenotypes.

## Acknowledgments

These studies were funded by US NIH grant R01 AI144126. We thank Tatiana Castro-Padovani and Jessamyn Moore for assistance and helpful comments on this manuscript.

---

**Supplemental Fig. 1.**
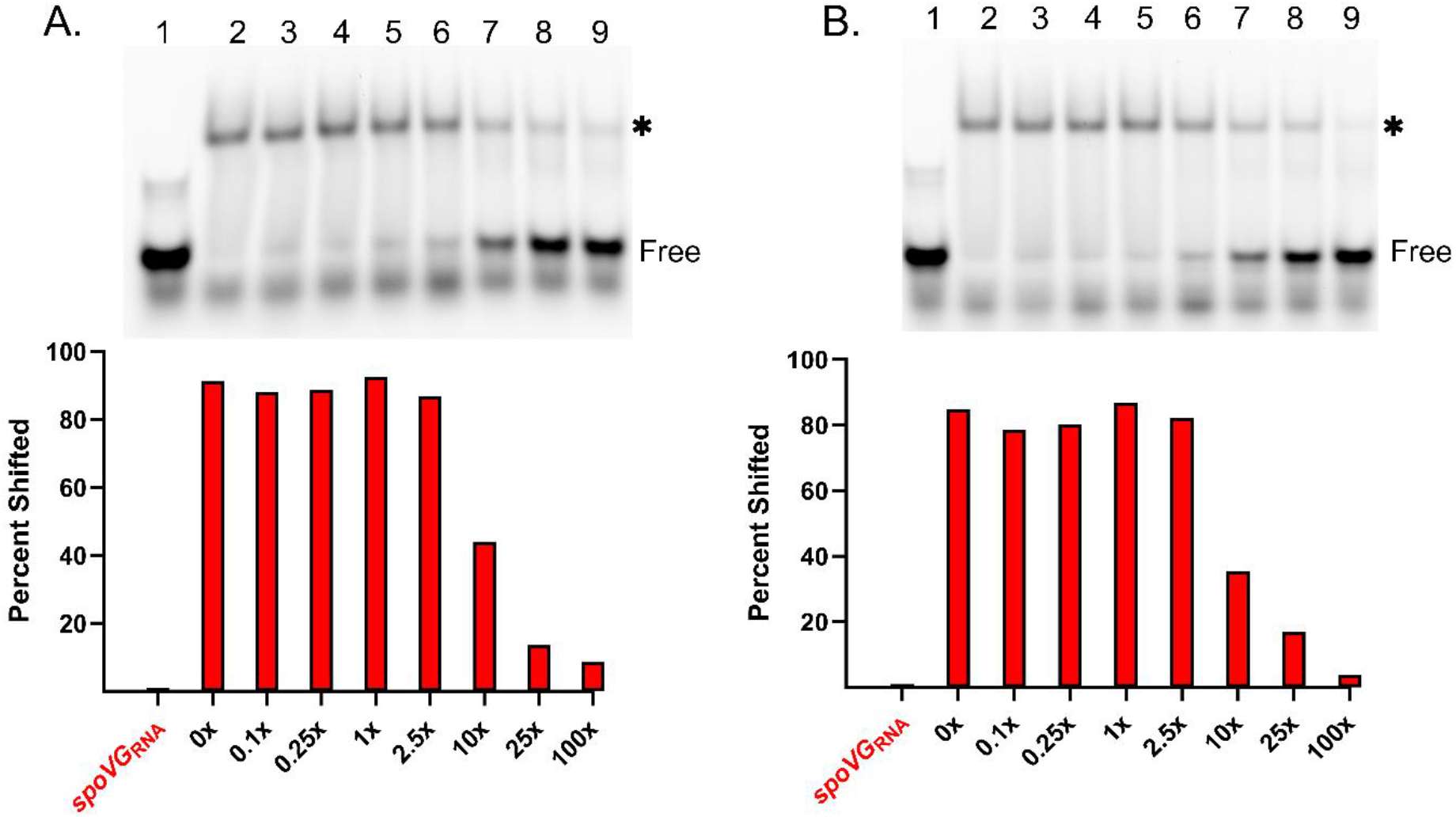
**A. and B.** Replicates of the competition assay using 100nM labeled *spoVG*_RNA_. Free probe is indicated, and the shifted probe is indicated by an asterisk. SpoVG concentration was 500nM in lanes 2-9. Unlabeled *spoVG*_RNA_ was added at 10nM, 25nM, 100nM, 250nM, 1μM, 25 μM, and 100 μM in lanes 3-9 respectively. EGTA was added to 5mM.

**Supplemental Fig. 2.**
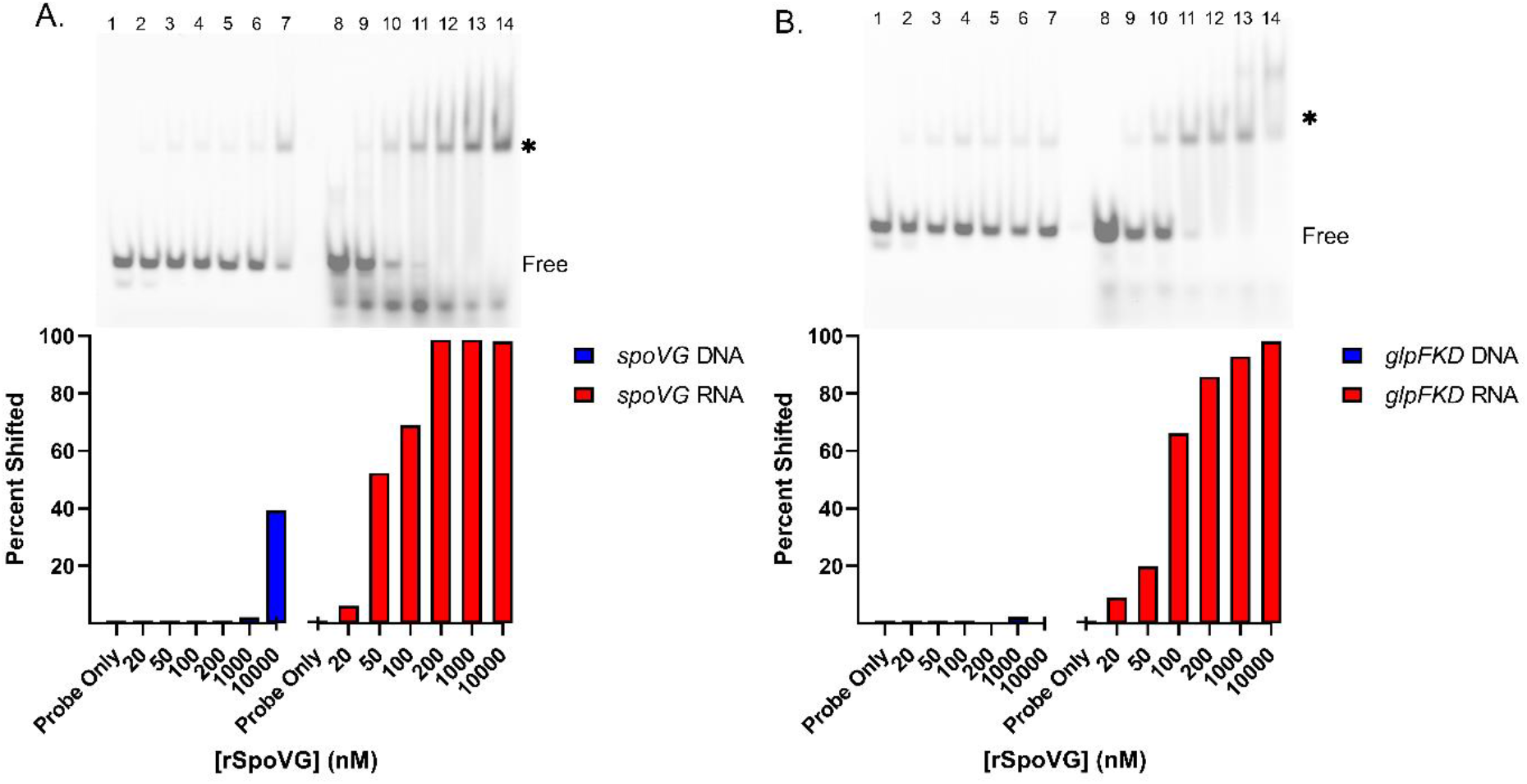
**A.** EMSA using labeled *spoVG*_DNA_ (lanes 1-7) and *spoVG*_RNA_ (lanes 8-14) at 100nM. Free probe is indicated, and the shifted probe is indicated by an asterisk. SpoVG was added to lanes 2-7 and 9-14 at 20nm, 50nm, 100nM, 200nM, 1μM, and 10μM. **B.** EMSA using labeled *glpFKD*_DNA_ (lanes 1-7) and *glpFKD*_RNA_ (lanes 8-14) at 100nM. SpoVG was added to lanes 2-7 and 9-14 at 20nm, 50nm, 100nM, 200nM, 1 μM, and 10μM.

**Supplemental Fig. 3.**
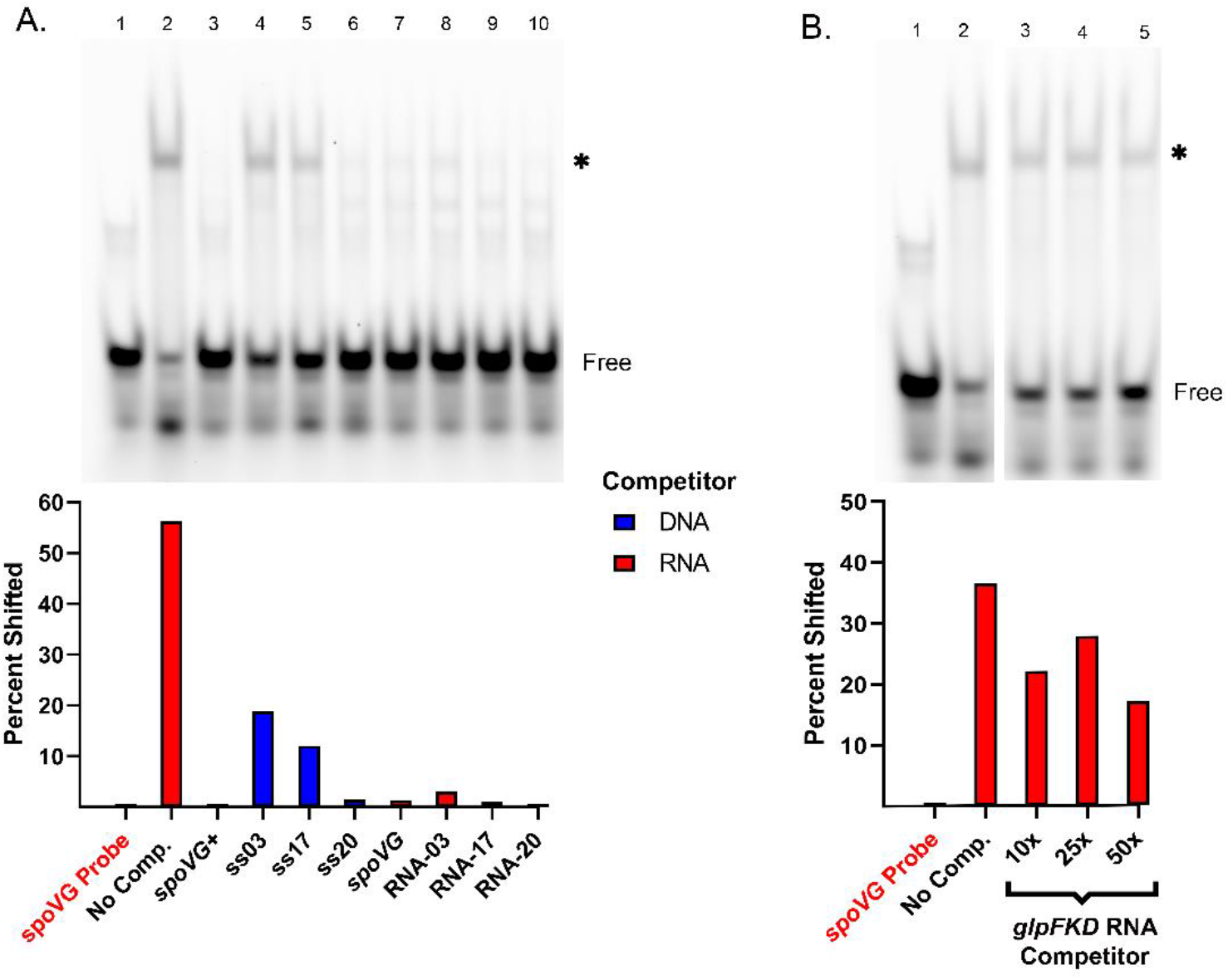
**A.** Competition assay using 100nM *spoVG*_RNA_ probe and 100nM of SpoVG in lanes 2-10. Free probe is indicated, and the shifted probe is indicated by an asterisk. Competitors were added at 5μM in the following sequence: Lane 3 *spoVG*_CD_. Lane 4 ss03; lane 5 ss17, lane 6 ss20, lane 7 unlabeled *spoVG*_RNA_; lane 8 RNA-03; lane 9 RNA-17; and lane 10 RNA-20. **B.** EMSA with 100nM of *spoVG*_RNA_ probe and 100nM of SpoVG added to lanes 2-5. Unlabeled *glpFKD*_RNA_ was added as competitor to lanes 3, 4, and 5 at 1μM, 2.5μM, and 5μM respectively.

**Supplemental Figure 4.**
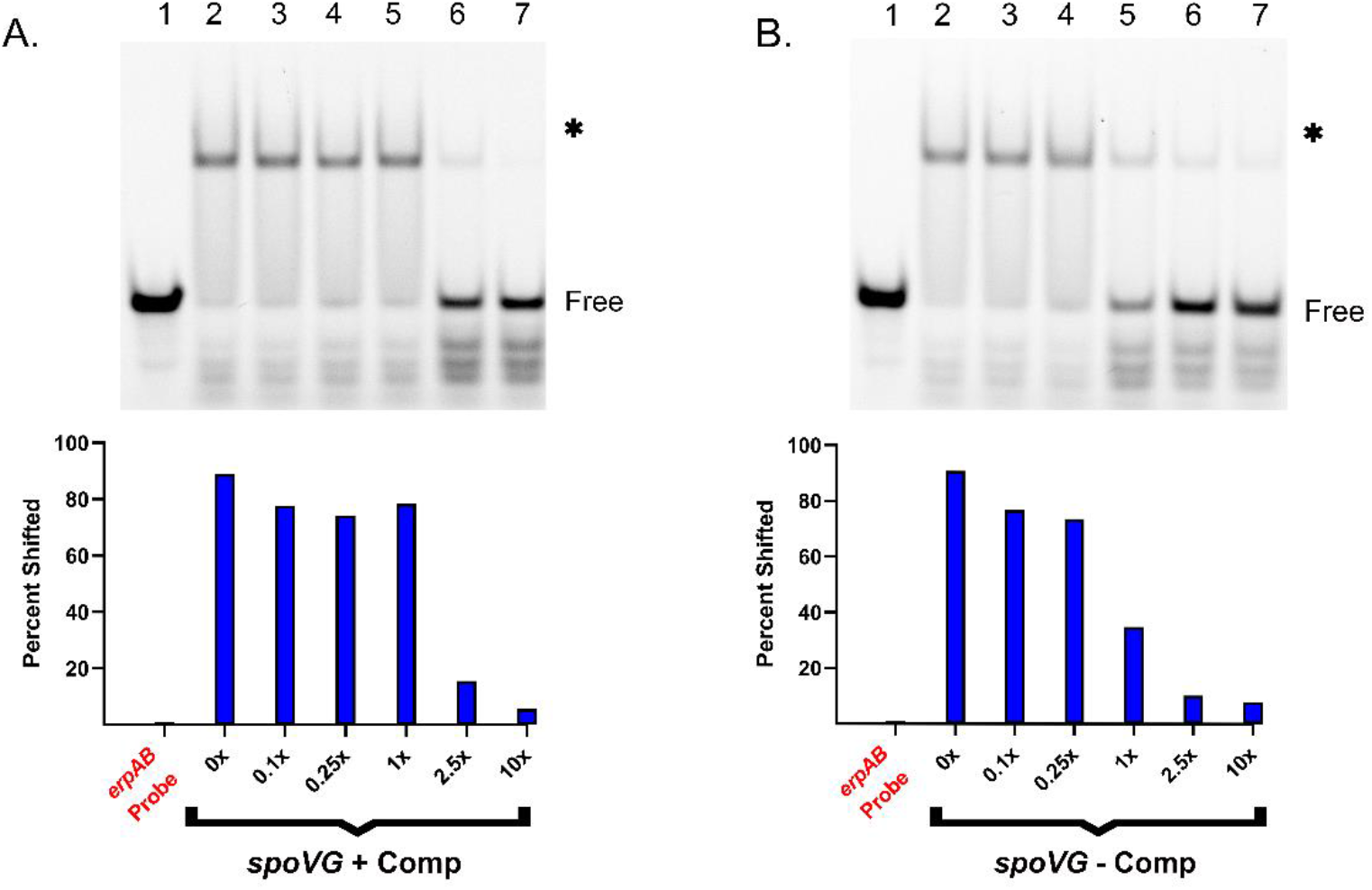
SpoVG binds differentially to the coding and non-coding strands of *spoVG*. **A.** EMSAs using 100nM of *erpAB*_RNA_ probe. SpoVG was added to 666 nM in lanes 2-7. Free probe is indicated, and the shifted probe is indicated by an asterisk. The SpoVG-*erpAB*_RNA_ complex was competed against using *spoVG*_CD_ in lanes 3-7 at 10nM, 25nM, 100nM, 250nM, 1μM, 2.5μM, and 10μM concentrations respectively. **B.** Repeat of EMSA in (A) except *spoVG*_NC_ ssDNA was used as the competitor.

**Supplemental Fig. 5.**
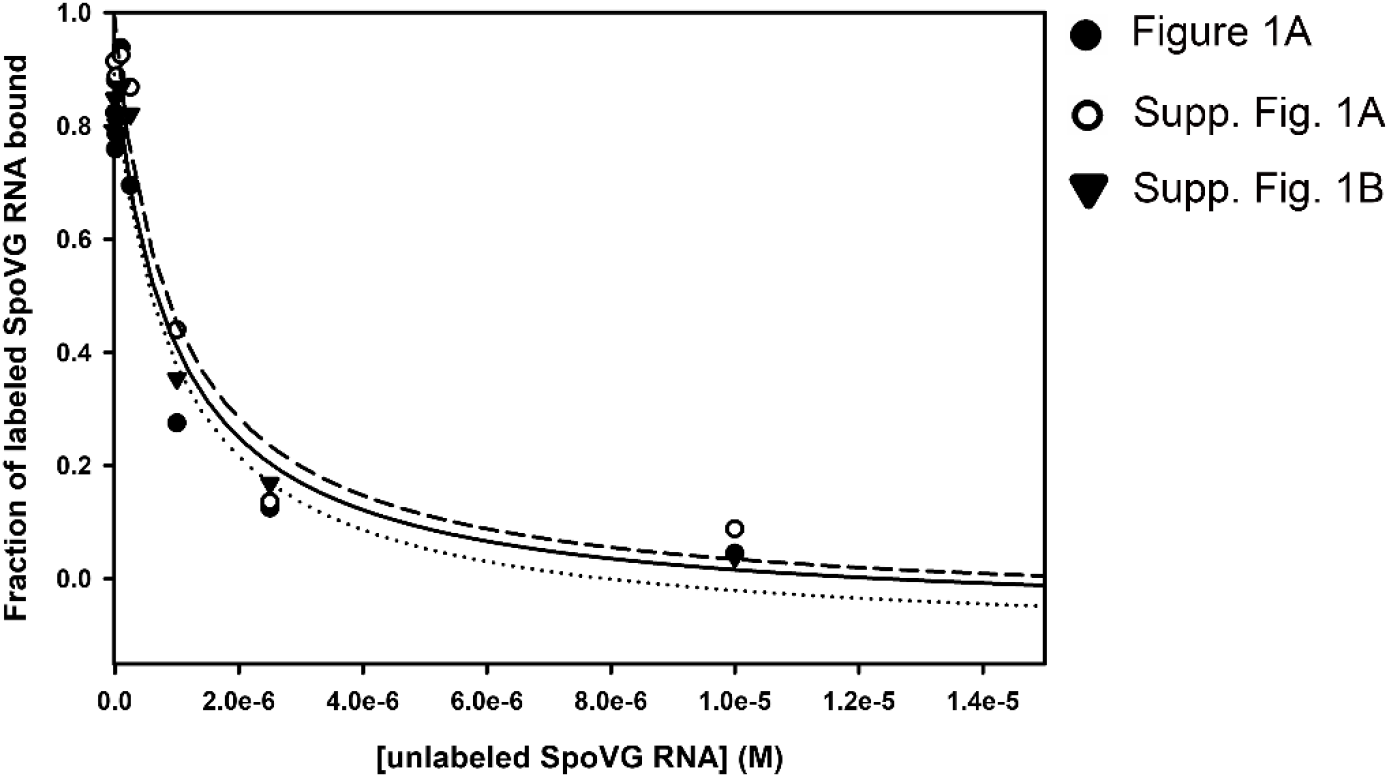
Graph of the K_D_ and IC50 calculations.

**Supplemental Table 1.**
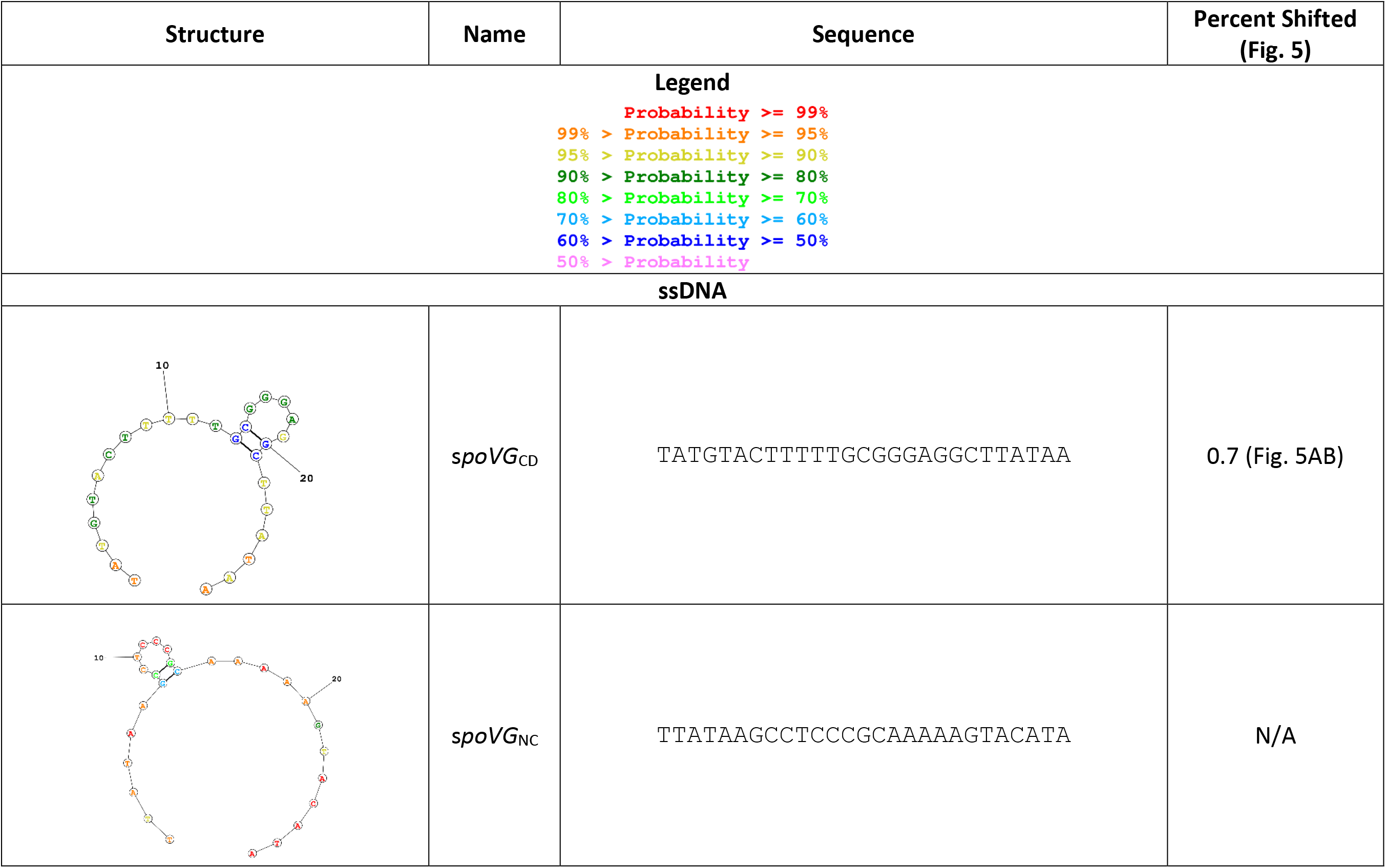
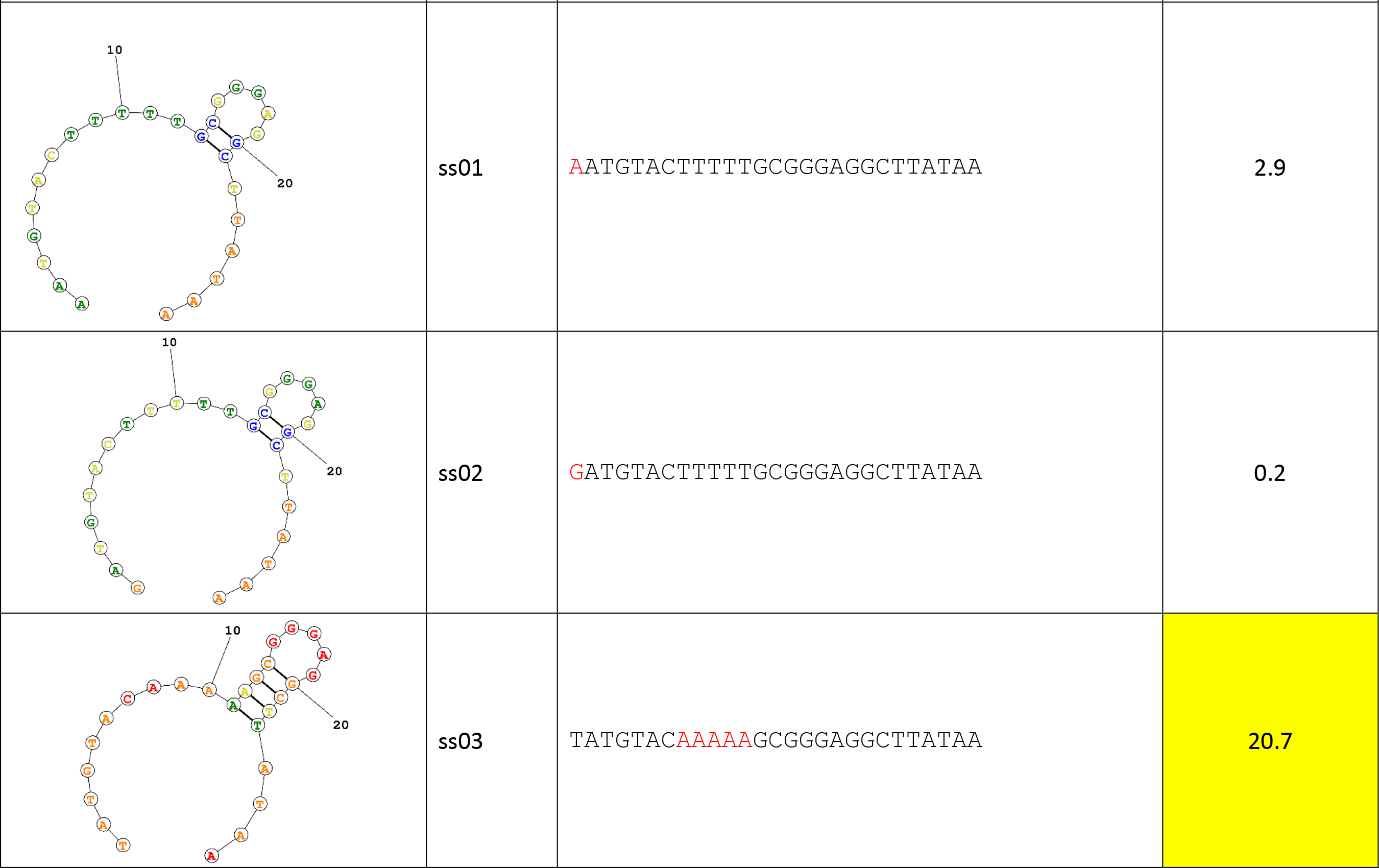
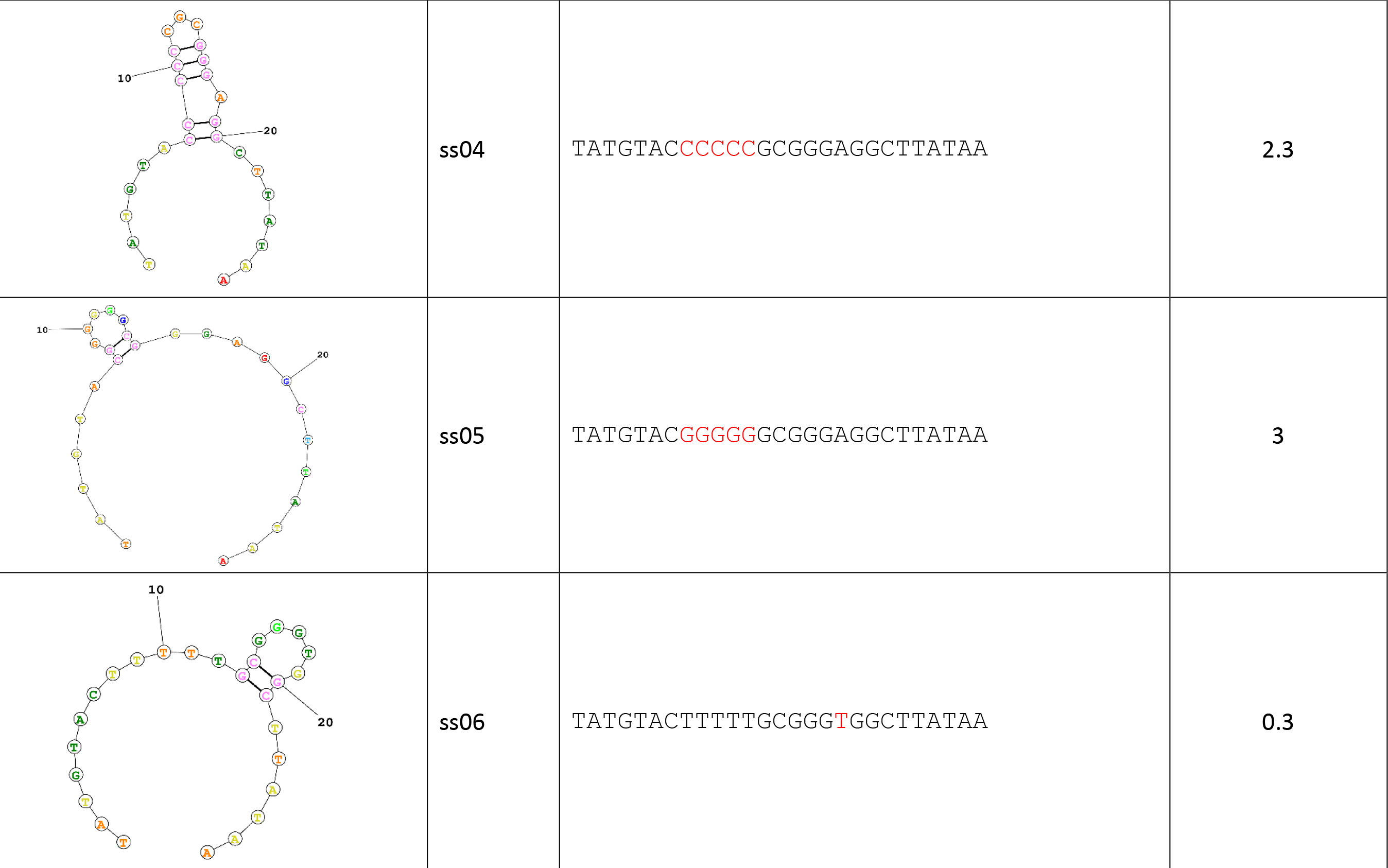
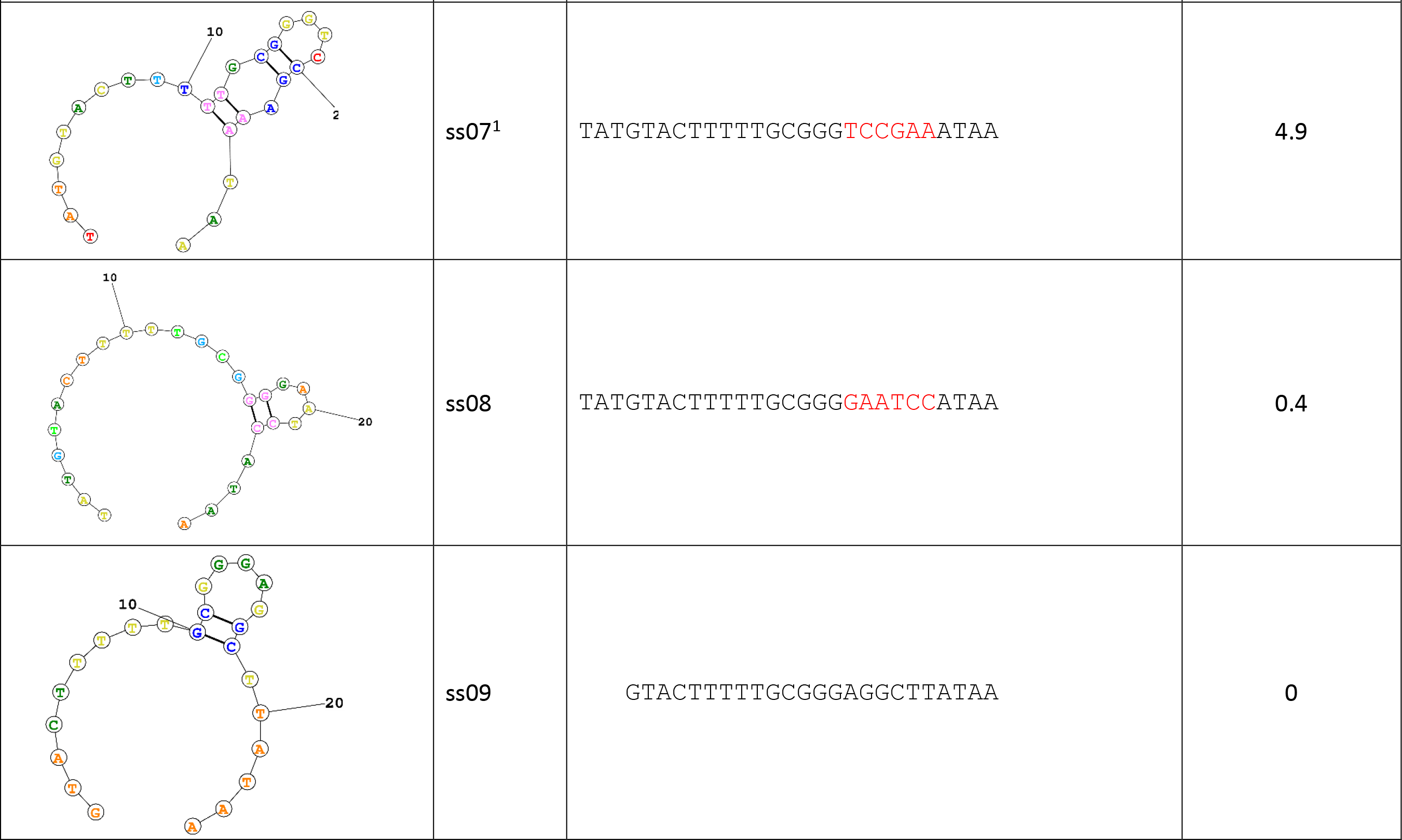
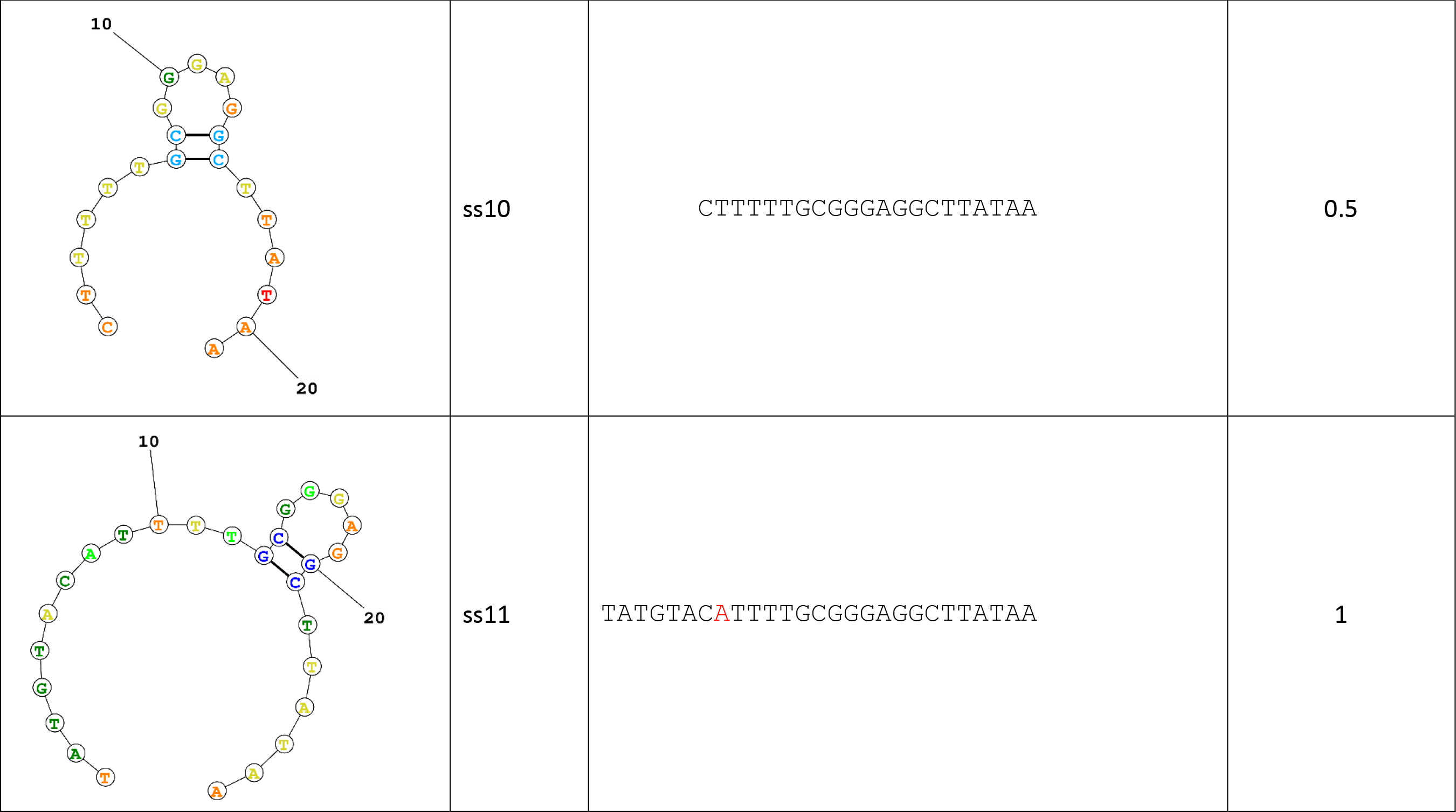
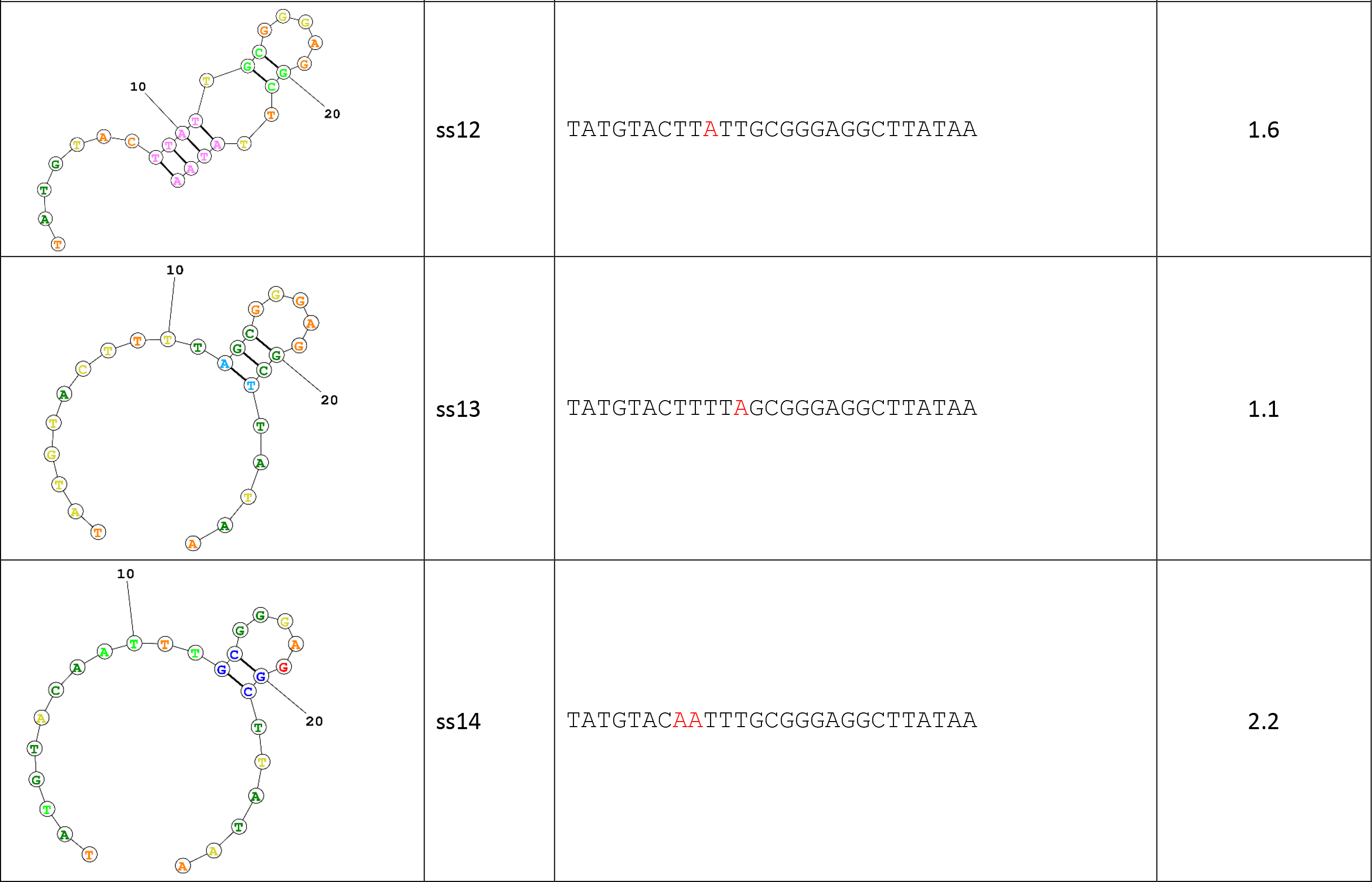
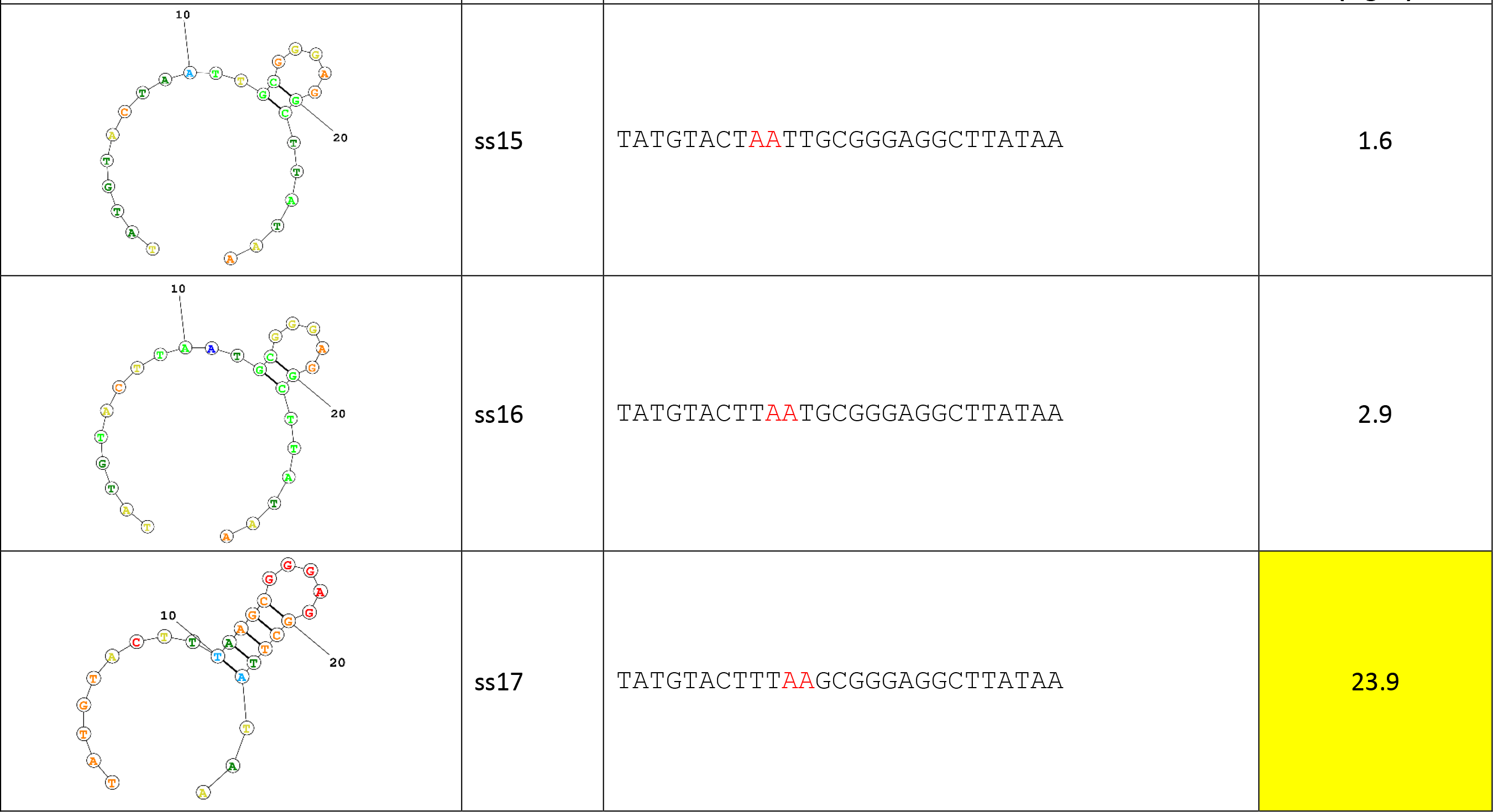
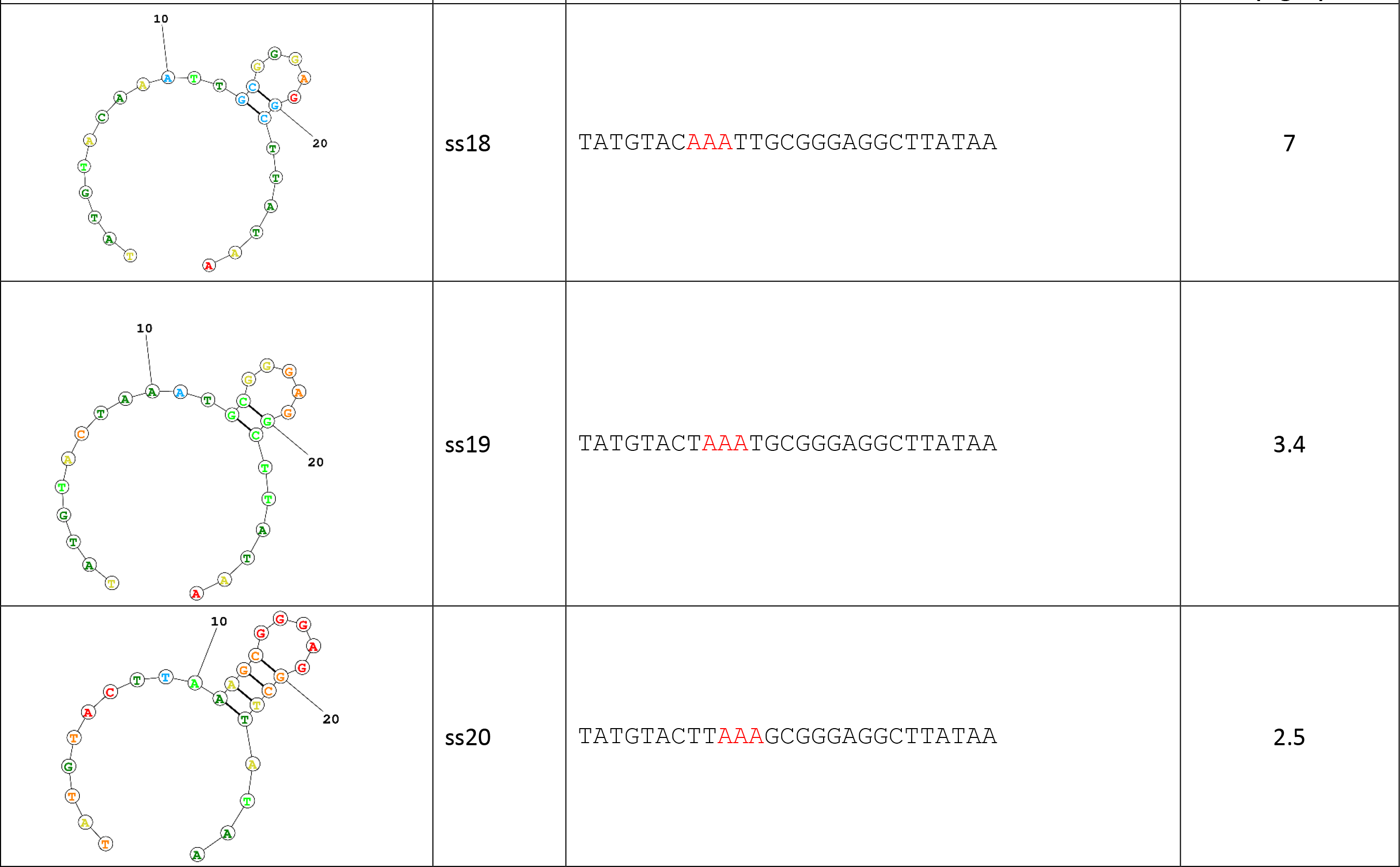
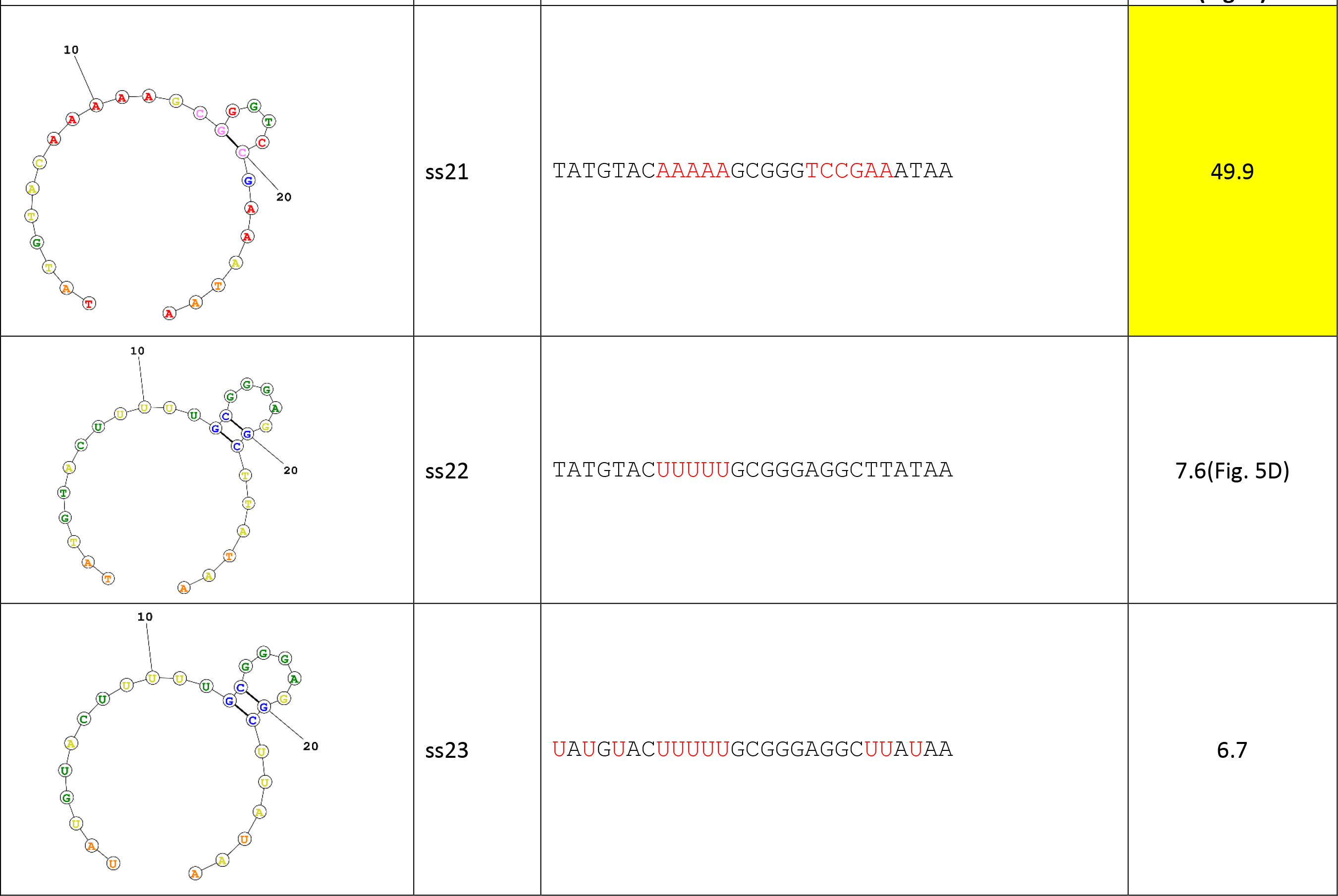
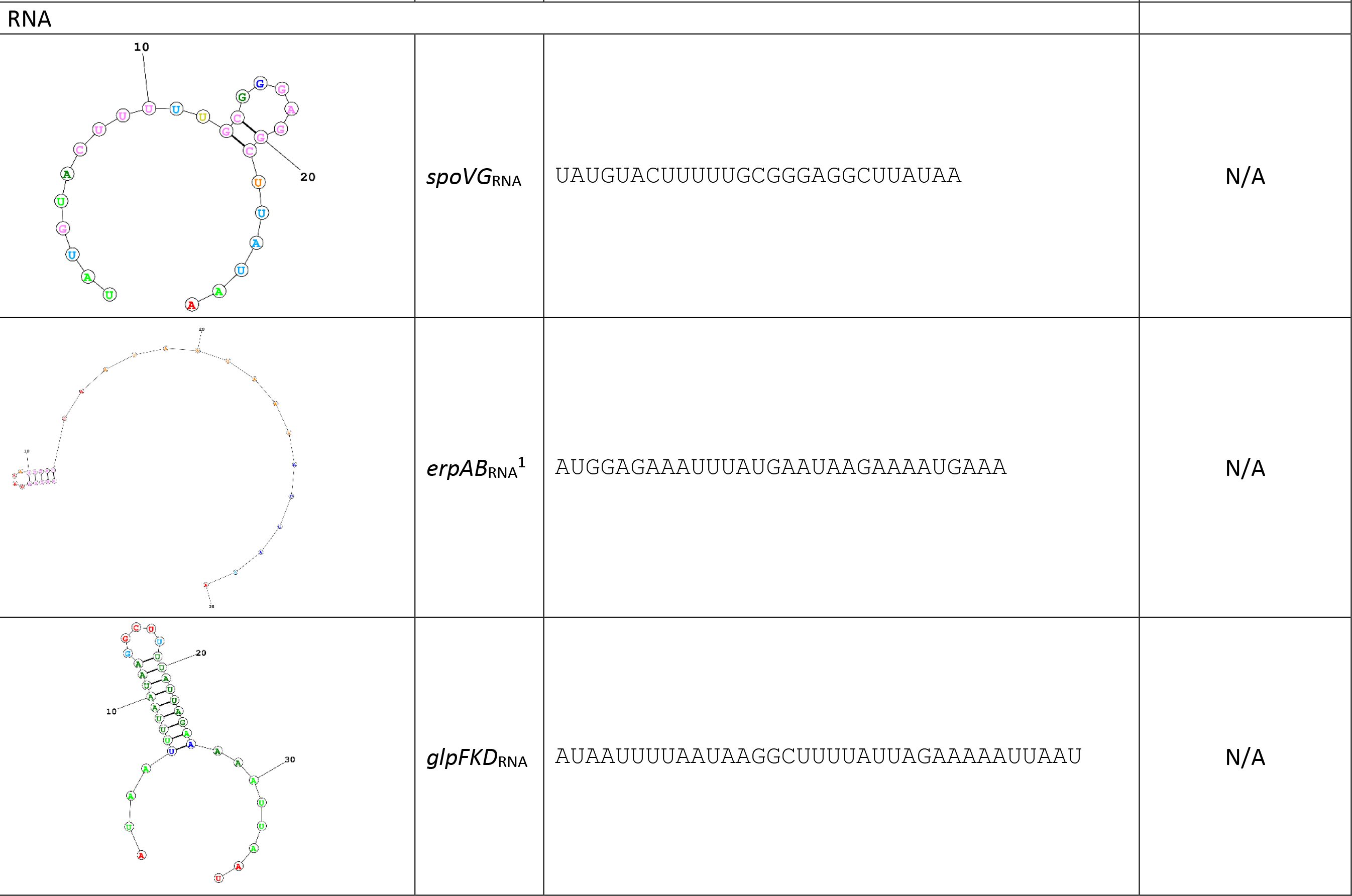
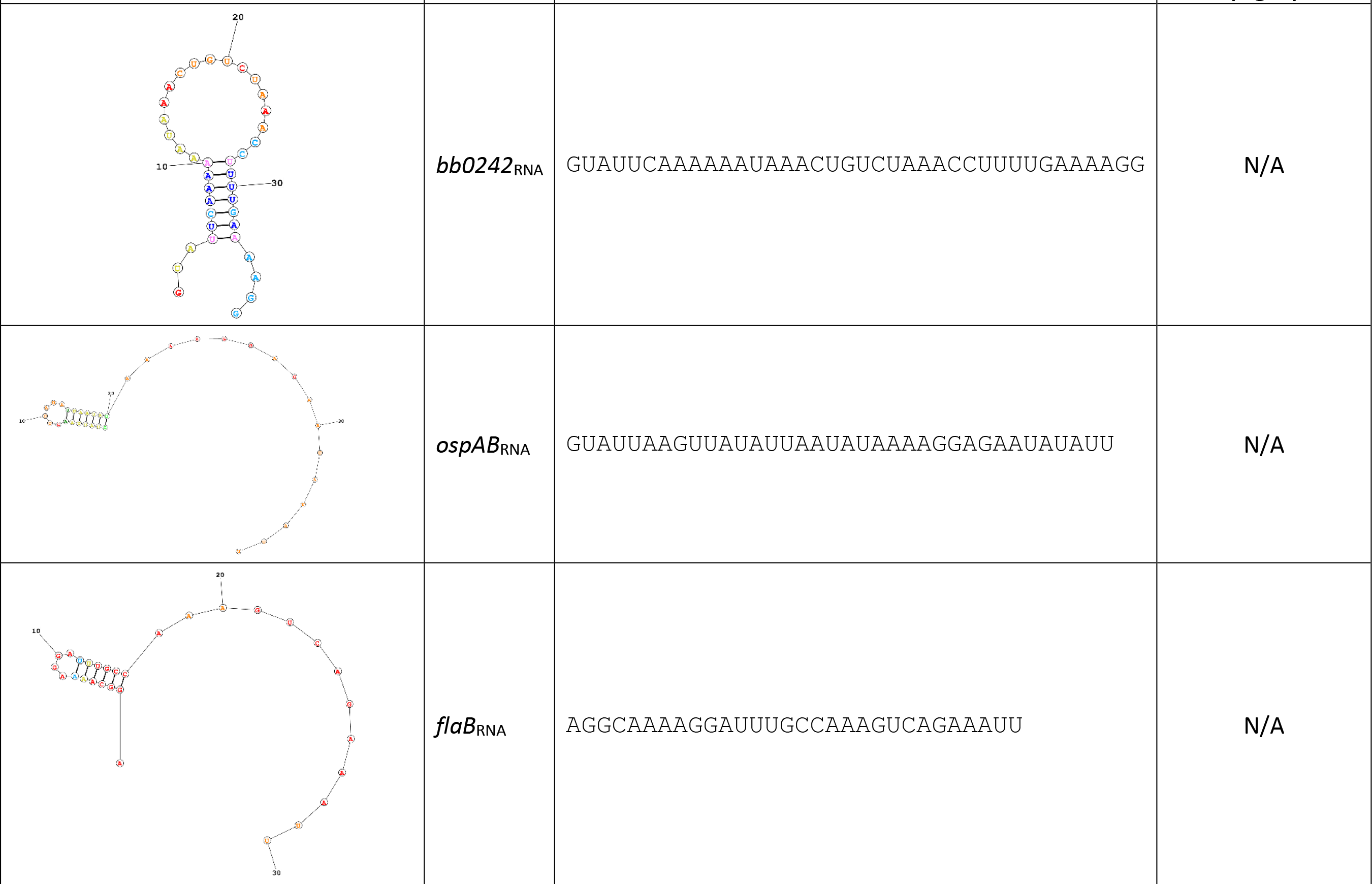
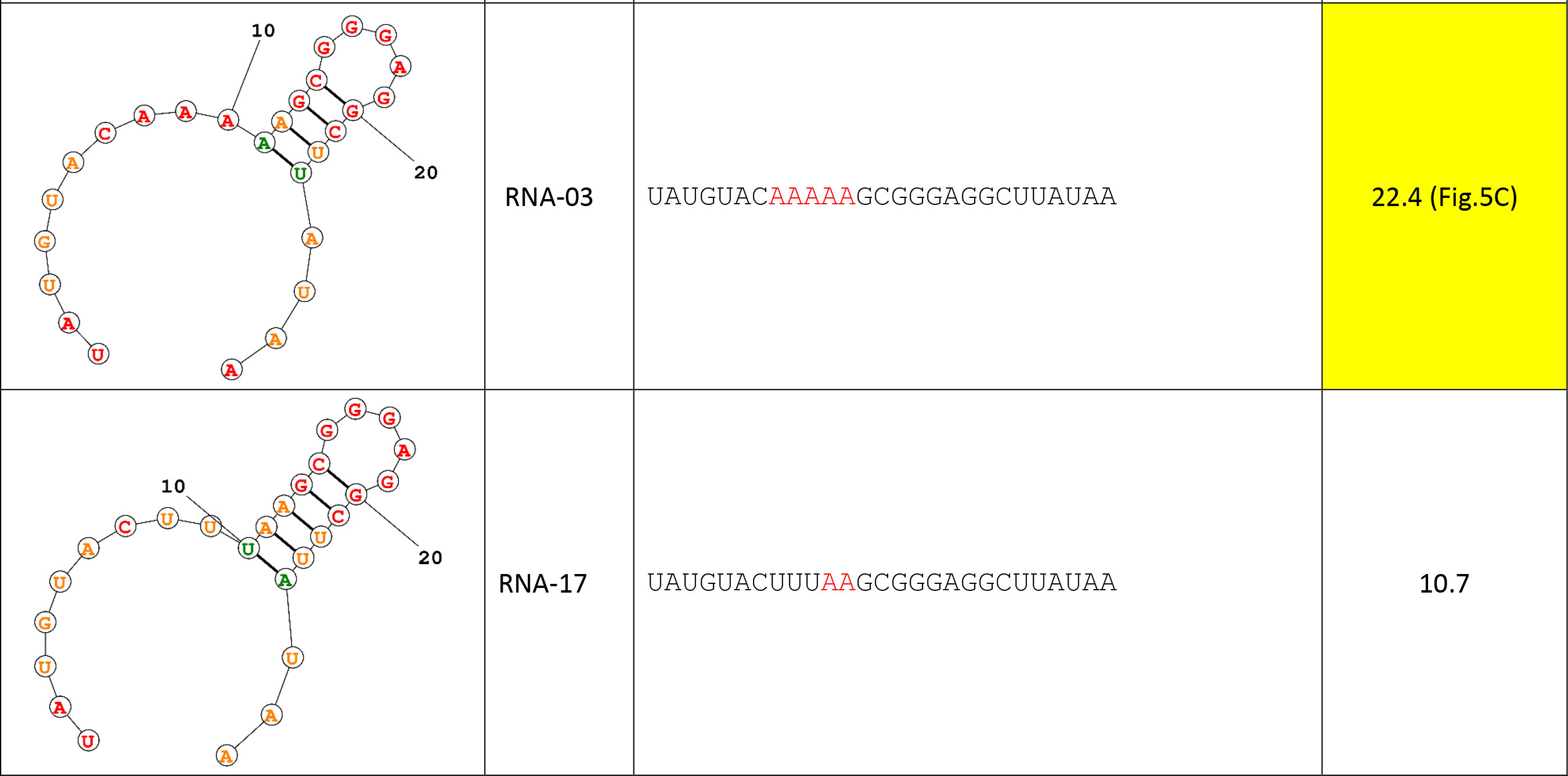
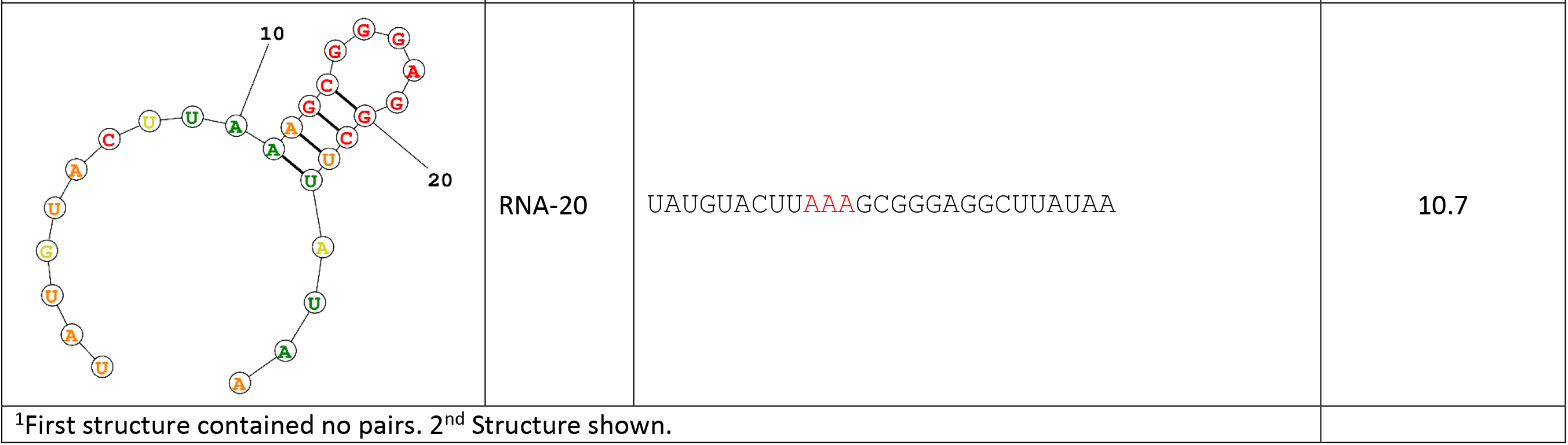
Predicted secondary structures of ssDNA and RNA oligonucleotides used. [19]

